# General anaesthesia reduces the uniqueness of brain connectivity across individuals and across species

**DOI:** 10.1101/2023.11.08.566332

**Authors:** Andrea I. Luppi, Daniel Golkowski, Andreas Ranft, Rudiger Ilg, Denis Jordan, Danilo Bzdok, Adrian M. Owen, Lorina Naci, Emmanuel A. Stamatakis, Enrico Amico, Bratislav Misic

**Affiliations:** Montréal Neurological Institute, McGill University, Montréal, QC, Canada; Department of Neurology, Klinikum rechts der Isar, Technical University Munich, Munich, Germany; Department of Anesthesiology and Intensive Care, School of Medicine and Health, Technical University of Munich, Munich, Germany; Asklepios Clinic, Department of Neurology, Bad Tolz, Germany; Department of Anaesthesiology and Intensive Care Medicine, Klinikum rechts der Isar, Technical University Munich, Munich, Germany; MILA, Quebec Artificial Intelligence Institute, Montréal, QC, Canada; Western Institute for Neuroscience (WIN), Western University, London, ON, Canada; Trinity College Institute of Neuroscience, School of Psychology, Trinity College Dublin, Dublin, Ireland; Division of Anaesthesia and Department of Clinical Neurosciences, University of Cambridge, Cambridge, UK; Neuro-X Institute, Ecole Polytechnique Federale de Lausanne, Lausanne, Switzerland

## Abstract

The human brain is characterised by idiosyncratic patterns of spontaneous thought, rendering each brain uniquely identifiable from its neural activity. However, deep general anaesthesia suppresses subjective experience. Does it also suppress what makes each brain unique? Here we used functional MRI under the effects of the general anaesthetics sevoflurane and propofol to determine whether anaesthetic-induced unconsciousness diminishes the uniqueness of the human brain: both with respect to the brains of other individuals, and the brains of another species. We report that under anaesthesia individual brains become less self-similar and less distinguishable from each other. Loss of distinctiveness is highly organised: it co-localises with the archetypal sensory-association axis, correlating with genetic and morphometric markers of phylogenetic differences between humans and other primates. This effect is more evident at greater anaesthetic depths, reproducible across sevoflurane and propofol, and reversed upon recovery. Providing convergent evidence, we show that under anaesthesia the functional connectivity of the human brain becomes more similar to the macaque brain. Finally, anaesthesia diminishes the match between spontaneous brain activity and meta-analytic brain patterns aggregated from the NeuroSynth engine. Collectively, the present results reveal that anaesthetised human brains are not only less distinguishable from each other, but also less distinguishable from the brains of other primates, with specifically human-expanded regions being the most affected by anaesthesia.

## INTRODUCTION

Consciousness—what is lost during anaesthesia and dreamless sleep, and restored upon awakening—is inherently subjective to each individual, as indicated by the near-synonymous use of “subjective experience” and “first-person experience”. In other words, each individual’s consciousness is unique to them. This raises an intriguing question: if consciousness is what makes each of us unique, do we become more alike when consciousness is lost?

A temporary state of unconsciousness can be induced by anaesthetics. The medical benefits of anaesthesia are well established, but anaesthesia is also finding increasing value as a tool to study the functioning of the brain [38, 39, 54, 55]. Unlike spontaneous sleep, anaesthetic-induced unconsciousness is amenable to experimental control: it can be reliably induced, maintained, and reversed.

Here we combine loss and recovery of consciousness induced by different anaesthetics, sevoflurane [29, 75] and propofol [46, 65], with functional MRI recordings of spontaneous activity in the human brain. We ask: does the human brain lose its distinctiveness when unconscious? We attack this question from three conceptual angles. First, we compare brains within and across individuals. Seminal work revealed that the patterns of functional connectivity (FC) between brain regions are reliably different across individuals, enabling “brain fingerprinting” of individuals based on fMRI scans [2, 24, 58, 88]. Therefore, here we use connectome fingerprinting to evaluate whether individuals become less distinguishable when under anaesthesia.

Second, we assess how well each individual’s brain activity across different levels of anaesthesia corresponds to canonical brain maps of cognitive operations obtained from meta-analytic aggregation of *>* 14, 000 neu-roimaging experiments [96]. Although our study concerns task-free fMRI, we reasoned that even in the absence of any tasks the brain may still spontaneously engage states pertaining to various cognitive operations [3, 18, 42, 60, 77, 80, 81]. In contrast, this should not occur during loss of consciousness, when even intrinsically-driven cognition should be abolished. This paradigm is inspired by evidence that the ability to detect brain responses to specific tasks (such as “imagine playing tennis” or “imagine navigating oround your house”) is a robust marker of consciousness even in individuals who are behaviourally unresponsive due to disorders of consciousness [5, 19–21, 36, 40, 61, 66–68].

Finally, we ask whether anaesthesia makes the human brain less distinctive from other species. Numerous reports show that under anaesthesia, the functional connectivity of different individuals regresses to a common template of structural connectivity [7, 22, 32, 51]. Motivated by this observation, we ask whether functional connectivity of the anaesthetised brain might also be understood as regressing towards a more fundamental template of primate functional connectivity—in other words, reducing the distinctiveness of our species compared to other nonhuman primates.

## RESULTS

Here, we consider resting-state functional MRI data obtained from N=15 healthy volunteers at baseline and after loss of behavioural responsiveness induced by different levels of the inhalational anaesthetic, sevoflurane: at EEG burst-suppression, 3 vol%, 2 vol%, and as well as during post-anaesthetic recovery of responsiveness [47, 75]. We replicate all results in an independent dataset of resting-state fMRI from N=16 healthy volunteers scanned before, during, and after loss of behavioural responsiveness induced by the intravenous anaesthetic, propofol [46, 65].

### Reduced identifiability of the anaesthetised brain

We first test the hypothesis that anaesthesia abolishes each individual’s idiosyncratic patterns of spontaneous neural activity, making the corresponding patterns of functional connectivity harder to distinguish across individuals. Specifically, correlate each individual’s FC during wakefulness with each individual’s FC during either post-anaesthetic recovery of responsiveness or anaesthesia. This produces an identifiability matrix where rows and columns are individuals, and each entry represents their connectomes’ similarity (correlation) (Fig. 1a).

**Figure 1.**
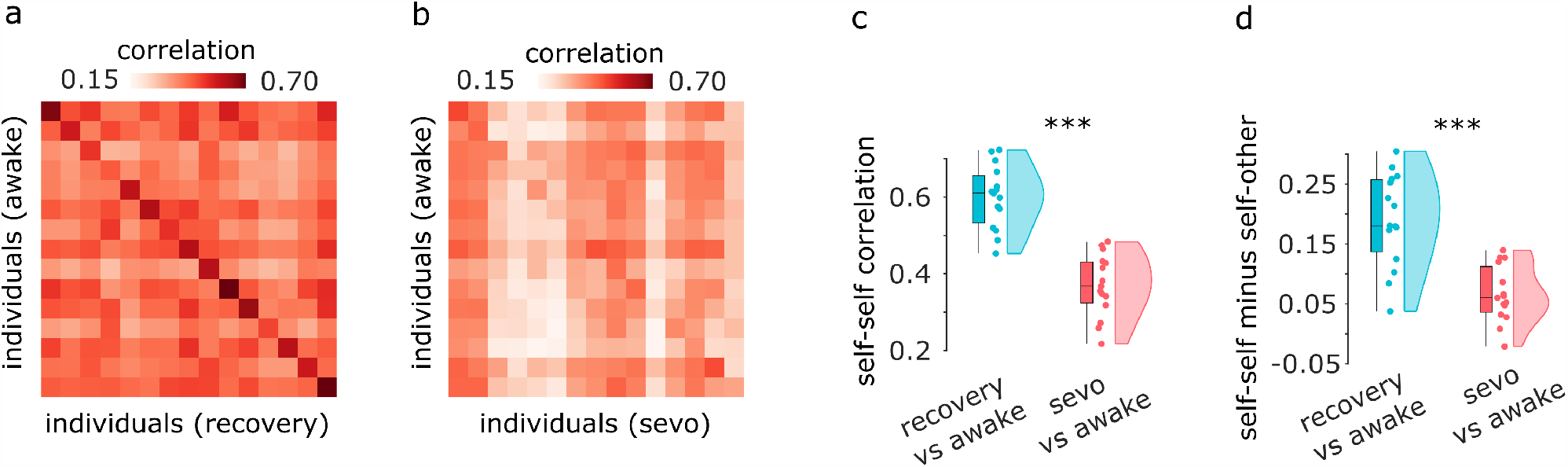
Identifiability of individual connectomes is diminished under sevoflurane anaesthesia. (**a**) Identifiability matrix between wakefulness and post-anaesthetic recovery. (**b**) Identifiability matrix between wakefulness and sevoflurane anaesthesia. Entries along the diagonal, represent self-self similarity (correlation of FC patterns), whereas off-diagonal entries represent self-other similarity. (**c**) Self-self similarity is significantly higher between two conscious states, than between wakefulness and anaesthesia. (**d**) Differential identifiability (the difference between self-self correlation and mean self-other correlation) is significantly higher between two conscious states, than between wakefulness and anaesthesia. Box-plots: center line, median; box limits, upper and lower quartiles; whiskers, 1.5*×* interquartile range. ***, *p <* 0.001.

We observe that when comparing two scans of the same individual while awake, it is easy to distinguish self from other. This identifiability can be discerned as a clear pattern along the matrix diagonal, representing self-self similarity, which is clearly distinguishable from off-diagonal entries, which represent self-other similarity (Fig. 1a). This is consistent with previous work on functional connectome fingerprinting using test-retest scans [2]. In contrast, the diagonal is barely discernible when awake scans are compared against anaesthetised ones, indicative of low identifiability of individuals (Fig. 1b). Indeed, self-self correlations between awake and anaesthetised brains are significantly diminished compared to awake-recovery self-similarity (awake-recovery: mean = 0.60, SD = 0.08; awake-anaesthesia: mean = 0.37; SD = 0.08; *t*(14) = 8.36, *p <* 0.001, effect size (Hedge’s *g*) = 2.71, CI [2.22, 3.69]; Fig. 1c). Likewise, differential identifiability (the difference between self-self correlation and mean self-other correlation) becomes significantly reduced when considering anaesthetised brains (awake-recovery: mean = 0.41, SD = 0.04; awake-anaesthesia: mean = 0.30; SD = 0.04; *t*(14) = 6.78, *p <* 0.001, effect size (Hedge’s *g*) = 2.50, CI [1.77, 3.85]; Fig. 1d). Collectively, these results demonstrate that anaesthetic-induced loss of responsiveness manifests as reduced distinctiveness of the individual functional connectome. We also show that analogous results are obtained at different depths of sevoflurane anaesthesia (Fig. S1). In the rest of this subsection, we identify the anatomical organization of functional connections that contribute to this change in distinctiveness.

We quantify edge-wise identifiability using the intraclass correlation coefficient (ICC), which describes how strongly elements in the same group resemble each other, for a given score. In this context, we obtain an ICC value for each edge (functional connectivity value between two brain regions), which indicates how well the weight of that edge separates within and between individuals [2]. Thus, the higher the ICC of an edge, the higher its identifiability (Methods) [2]. The difference in edgewise identifiability between awake-recovery and awake-anaesthesia therefore indicates the extent to which identifiability of each functional edge is affected by anaesthesia. We find that sevoflurane anaesthesia reduces the contribution to identifiability of virtually all edges (Fig. 2a). This pattern is neither uniform nor random: rather, functional connections that most diminish their contribution to identifiability are those with an underlying direct structural connection, as quantified using in vivo diffusion MRI tractography (connected: mean = 0.34, SD = 0.35; disconnected: mean = 0.27; SD = 0.36; *t*(39998) = 12.96, *p <* 0.001, effect size (Hedge’s *g*) = 0.19, CI [0.16, 0.22]; Fig. 2b). Thus, functional connections between structurally connected regions are particularly vulnerable to anaesthesia, losing their distinctiveness and becoming more similar across individuals.

**Figure 2.**
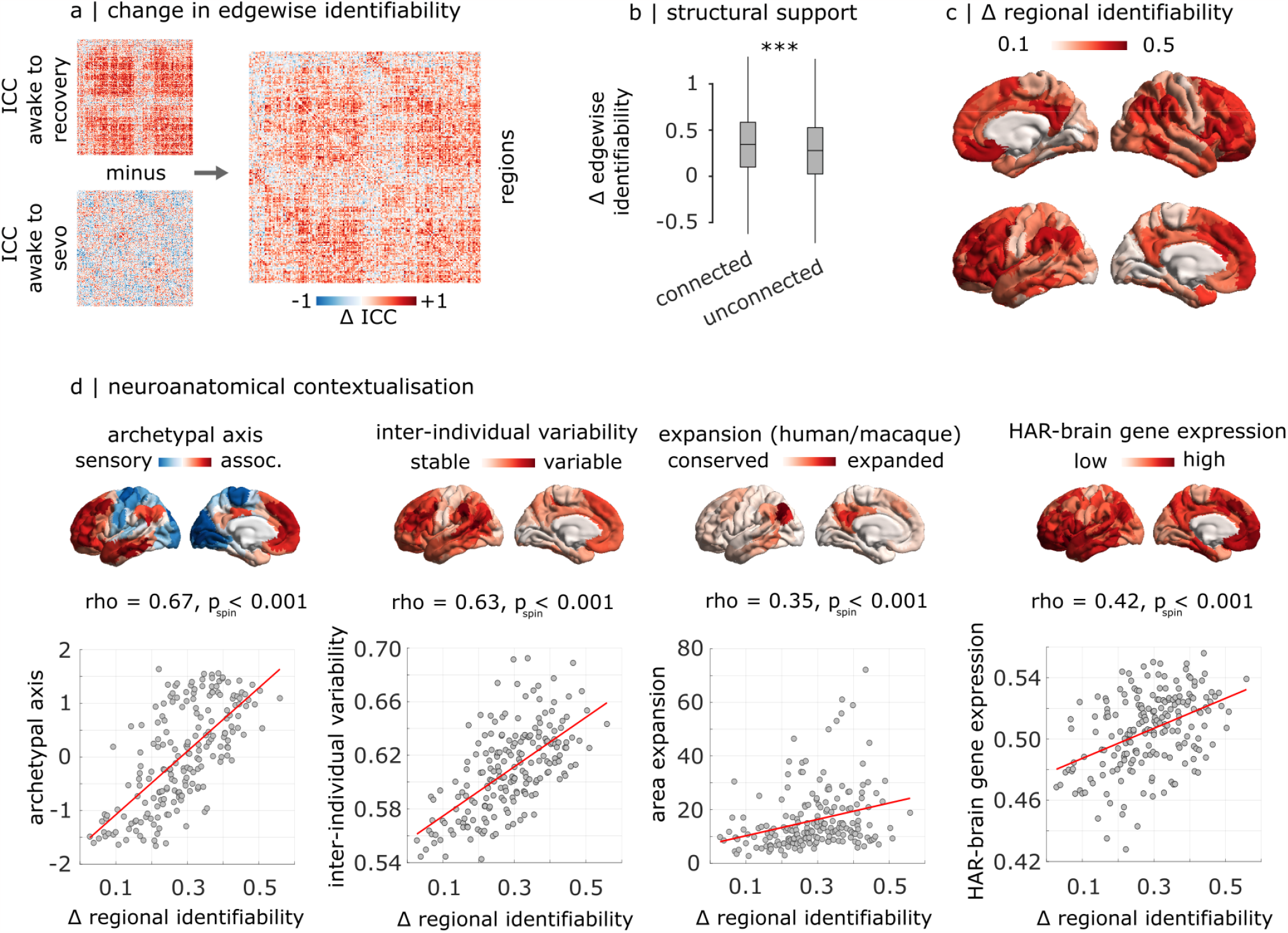
Anatomical characterisation of contributions to sevoflurane-induced loss of identifiability. (**a**) Edge-level difference in intra-class correlation coefficient between awake-recovery and awake-sevoflurane. (**b**) The anaesthetic-induced loss of ICC is significantly more pronounced for functional connections with an underlying structural direct connection, than without. ***, *p <* 0.001. (**c**) Regional distribution of anaesthetic-induced loss of ICC, projected onto the cortical surface. (**d**) The anaestheticinduced regional loss of ICC is significantly spatially aligned with the archetypal sensory-association axis of cortical organisation; the regional distribution of inter-individual variability of functional connectivity; the regional distribution of cortical expansion between macaque and human brains; and the regional expression of human-accelerated genes pertaining to brain function and development (“HAR-brain genes”).

We next localise regional changes in identifiability, quantified as the mean change in edgewise identifiability across each region’s edges. The greatest decreases in identifiability occur in regions of the default mode and fronto-parietal networks, and transmodal cortex more broadly, whereas unimodal (somatomotor and visual) cortices are least affected (Fig. 2c and Fig. S2). This pattern of unimodal-transmodal distinction is confirmed by a significant spatial correlation with the brain’s archetypal sensory-association axis (*ρ* = 0.67, *p*_*spin*_ *<* 0.001, *N* = 200 regions; Fig. 2d) [83]. Given that functional connectivity of transmodal cortex exhibits the greatest inter-individual variability in the awake state [62], we sought to test whether anaesthesia-induced reduction in identifiability preferentially targets these regions. Indeed, patterns of reduced identifiability are correlated with the map of inter-individual variability developed by Mueller and colleagues [62] (*ρ* = 0.63, *p*_*spin*_ *<* 0.001, *N* = 200 regions; Fig. 2d).

Having observed that changes in regional identifiability are more pronounced where individuals most differ in terms of FC, we further investigate whether a more general phenomenon is at play. Do anesthetic-induced changes in regional identifiability reflect not only distinctiveness between individuals, but more generally distinctiveness between species? Although we pursue this in more detail in a subsequent section, here we show that changes in regional identifiability correlate with molecular and morphometric markers of phylogenetic cortical differentiation between human and non-human primates. Specifically, anaesthetic-induced changes in regional identifiability are spatially correlated with the cortical map of evolutionary expansion between macaque and human [94] (*ρ* = 0.35, *p*_*spin*_ *<* 0.001, *N* = 200 regions; Fig. 2d), with greater change in distinctiveness being observed in phylogenetically newer regions. Likewise, we observe a significant spatial correlation between the regional changes in identifiability, and the regional mean expression of human-accelerated genes pertaining to brain function and development (“HAR-brain genes”; *ρ* = 0.42, *p*_*spin*_ *<* 0.001, *N* = 200 regions; Fig. 2d). These are genes associated with loci that displayed accelerated divergence in the human lineage compared with the chimpanzee, and therefore indicate evolution-related changes in the corresponding regions [92]. In other words, regions that exhibit greater change in distinctiveness, also exhibit greater expression of human-accelerated genes. Altogether, anaesthesia selectively reduces the identifiability of regions that are most distinctive, both between individuals, and between humans and non-human primates.

### Reduced match between spontaneous activity and canonical cognitive maps under anaesthesia

An influential paradigm to investigate pathological or pharmacological perturbations of consciousness is to determine whether cognitive processes can be algorithmically inferred (or ‘decoded’) from neural activity. For example, patients suffering from disorders of consciousness may be asked to imagine playing tennis while in the scanner, to determine whether motion-related regions become reliably activated in response to command, despite the absence of overt behavioural command-following [61, 68]). Although this is typically done in the presence of explicit tasks or other stimuli (e.g., movie-watching [61, 67, 68]), here we sought to investigate whether covert cognitive processes can be discerned from spontaneous neural activity more generally, through a comprehensive assay based on meta-analytic maps from thousands of neuroimaging experiments [31, 96]. Specifically, across different levels of anaesthesia we assess how well each individual’s neural activity maps at each point in time correspond to 123 canonical brain maps obtained from meta-analytic aggregation of *>* 14, 000 neuroimaging experiments [96] (Fig. 3a). For simplicity, we hereafter refer to this activity-based reverse inference approach as “cognitive matching”. Averaging across the scan duration provides, for each individual and each condition, an overall index of the quality of cognitive matching.

**Figure 3.**
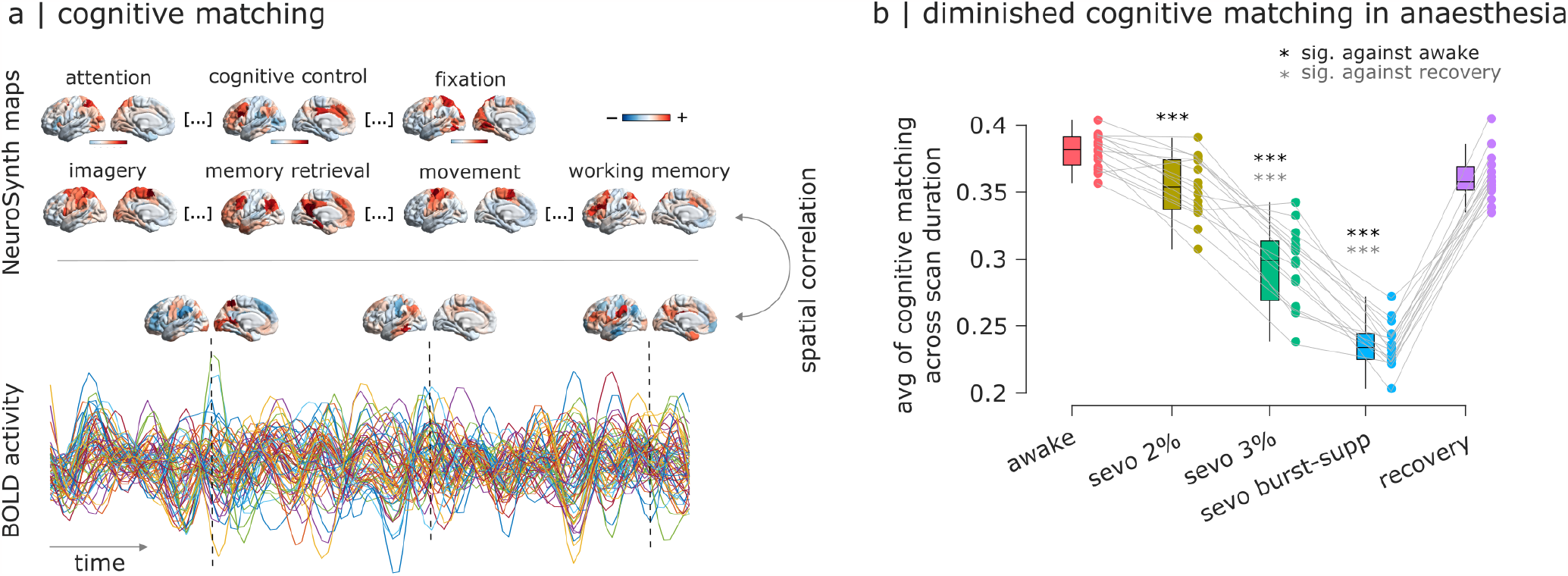
Cognitive matching of brain activity with canonical meta-analytic patterns is diminished under anaesthesia and restored upon recovery. (**a**) At each point in time, the cognitive matching score is computed as the best spatial correlation between brain activity and 123 NeuroSynth meta-analytic maps. For each participant, an overall index of the quality of cognitive matching is then obtained by averaging the cognitive matching scores across the entire scan duration, within each condition. (**b**) Ordinate: mean across time of the cognitive matching score. *** (black), *p <* 0.001 against wakefulness (FDR-corrected); *** (gray), *p <* 0.001 against recovery (FDR-corrected). Box-plot: center line, median; box limits, upper and lower quartiles; whiskers, 1.5*×* interquartile range.

We find that as anaesthesia deepens (increasing concentration of sevoflurane), the quality of cognitive matching deteriorates: the best spatial correlation between brain activity and meta-analytic brain maps from NeuroSynth (averaged across the entire scan duration) is lower at deeper levels of anaesthesia (Fig. 3b). This trend is reversed upon recovery of responsiveness (Fig. 3b; full statistics shown in Table S1). The anaesthetic-induced reduction in the quality of cognitive matching is more pronounced for NeuroSynth maps that load onto the higher-order (transmodal/association) end of the brain’s archetypal axis (e.g. “cognitive control”, “emotion regulation”), than for maps that load onto the unimodal/sensory end (e.g. “fixation”, “movement”; Fig. S3). In other words, anaesthesia diminishes the extent to which spontaneous brain activity reflects cognitive patterns from the literature, particularly for higher-order cognitive operations, potentially explaining why individual distinctiveness is suppressed by anaesthesia.

### Anaesthesia increases similarity of functional connectivity between humans and non-human primates

Finally, we investigate whether there is greater similarity between the patterns of human and macaque functional connectivity when humans are awake, or when they are anaesthetised. We used functional MRI data from macaques scanned while awake [57], and processed similarly to human data [48]. The human data were in turn re-parcellated according to the Regional Mapping parcellation of Kötter and Wanke [41], translated between macaque and human brains by [12], such that each cortical region is anatomically matched to its homologue across the two species. Upon comparing the correlation between human and macaque patterns of cortical functional connectivity, we find that indeed, the deepest levels of sevoflurane induce significantly greater similarity between the two species, going from predominantly negative correlations to fully positive ones (Fig. 4). Once again, we also see that this effect is reversed upon recovery of responsiveness (Fig. 4 and Table S2). Thus, anaesthesia increases the similarity between the functional connectivity of the human brain, and the functional connectivity of the non-human primate brain. This result complements our observation that anaesthetic-induced reduction in regional identifiability is most pronounced in regions of the human brain that are genetically most human-specific (Fig. 2d).

**Figure 4.**
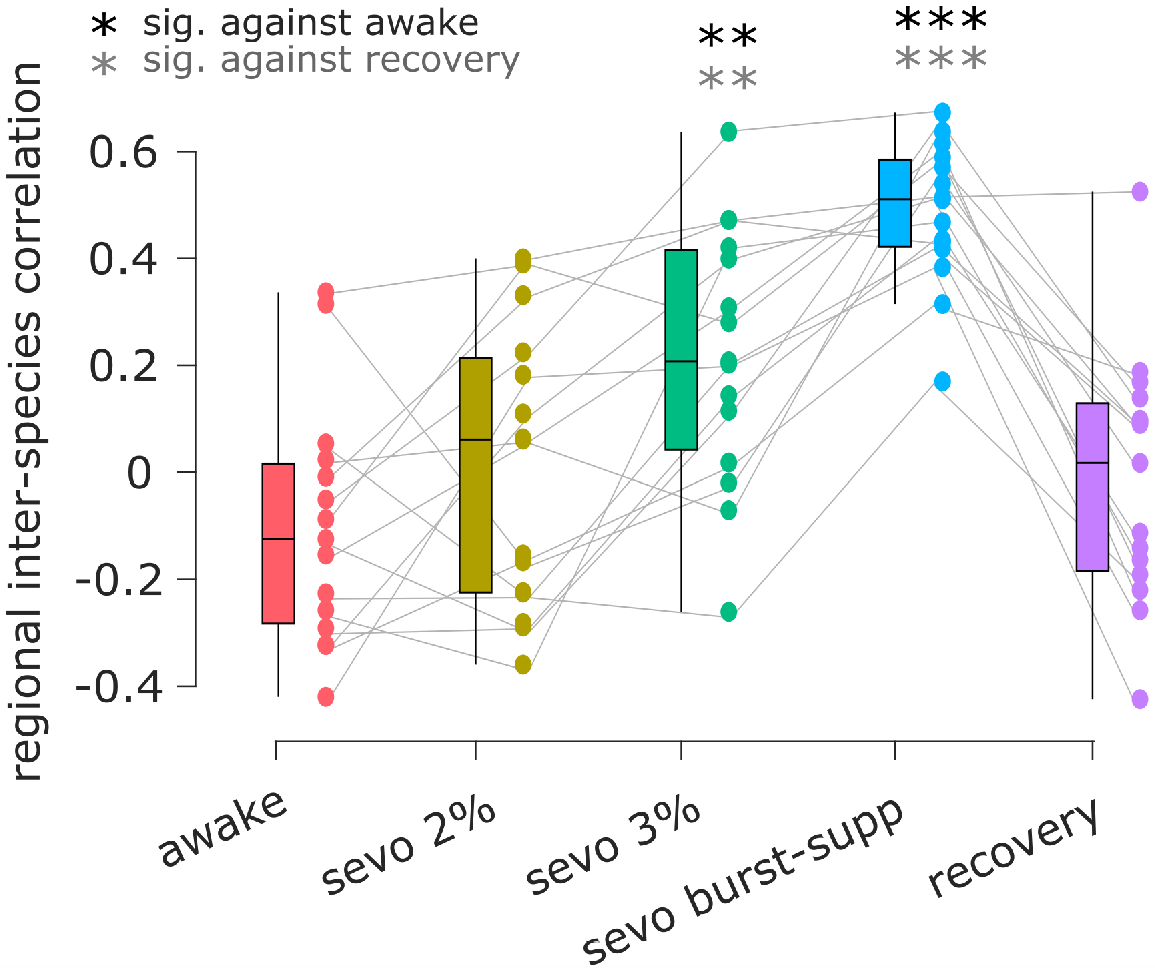
Human cortical functional connectivity is more similar to macaque cortical connectivity under anaesthesia than during wakefulness. Ordinate: Regional correlation of mean FC between human and macaque across wakefulness, anaesthesia, and recovery. **, *p <* 0.01, *** (black), *p <* 0.001 against wakefulness (FDR-corrected); ** (gray), *p <* 0.01 *** (gray), *p <* 0.001 against recovery (FDR-corrected). Box-plots: center line, median; box limits, upper and lower quartiles; whiskers, 1.5*×*interquartile range.

### Replication and robustness

#### Replication with propofol anaesthesia

We replicate all results in a separate dataset of anaesthesia with the intravenous agent, propofol [46, 65]. Although this dataset does not reach the same depth of anaesthesia as used in the main analysis, nevertheless, the results are broadly consistent with what is observed under sevoflurane anaesthesia. Self-self similarity decreases during anaesthetic-induced loss of behavioural responsiveness (awake-recovery: mean = 0.68, SD = 0.07; awake-anaesthesia: mean = 0.48; SD = 0.13; *t*(15) = 6.81, *p <* 0.001, effect size (Hedge’s *g*) = 1.82, CI [1.32, 2.74]) and so does differential identifiability (awake-recovery: mean = 0.20, SD = 0.06; awake-anaesthesia: mean = 0.08; SD = 0.05; *t*(15) = 7.19, *p <* 0.001, effect size (Hedge’s *g*) = 2.03, CI [1.41, 3.01]; Fig. S4). Likewise, the regional distribution of propofol-induced changes in identifiability is also more pronounced in default mode and fronto-parietal than somatomotor and visual cortices, correlating with the sensory-association axis (*ρ* = 0.46, *p*_*spin*_ *<* 0.001, *N* = 200 regions). We also replicate the correlation between the regional change in identifiability, and canonical maps of inter-individual variability (*ρ* = 0.43, *p*_*spin*_ *<* 0.001, *N* = 200 regions), evolutionary cortical expansion (*ρ* = 0.28, *p*_*spin*_ = 0.001, *N* = 200 regions) and expression of HAR-brain genes (*ρ* = 0.30, *p*_*spin*_ *<* 0.001, *N* = 200 regions) (Fig. S5).

The cognitive matching results from NeuroSynth do not indicate statistically significant differences between propofol anaesthesia and either baseline or postanaesthetic recovery of consciousness, in terms of the maximal observed correlation between brain activity and meta-analytic maps (Fig. S6 and Table S3). However, if instead of only considering the best correlation, we consider the average magnitude of correlations between brain activity and all NeuroSynth maps, then we do find significant differences between baseline and anaesthesia, both in the sevoflurane and propofol datasets (Fig. S6). This latter analysis may be interpreted as the overall ability of meta-analytic patterns to recapitulate patterns of spontaneous brain activity. Additionally, we also observe that propofol significantly increases the similarity between human and macaque regional FC, an effect that is then reversed upon recovery (Fig. S8 and Table S4).

#### Replication of cognitive matching with with BrainMap

The results pertaining to quality of decoding of brain activity based on the NeuroSynth meta-analytic engine are also replicated using 66 unique behavioural domains obtained from an alternative meta-analytic database, BrainMap [27, 43]. Whereas NeuroSynth has a data-driven bottom-up approach to taxonomy, and uses an automated process to identify statistical associations between brain coordinates and studies involving specific cognitive and behavioural terms, BrainMap is expert-curated. Despite the differences between the two databases (e.g. BrainMap explicitly excludes patient studies), we still find that the quality of decoding significantly deteriorates as the level of anaesthesia deepens, and is restored upon recovery of responsiveness (Fig. S9 and Table S6).

#### Robustness to choice of parcellation

We show that the present results – obtained using a functional atlas – can be replicated when using an alternative parcellation of the cerebral cortex, the Desikan-Killiany anatomical atlas [23] (Fig. S10 and Table S5). Likewise, similar results are obtained when including 32 subcortical regions as defined by the recent Tian atlas (Fig. S11 and Table S7). In particular, among subcortical structures we observe especially high regional contribution of the bilateral globus pallidus to the anaesthetic-induced change in regional identifiability.

#### Robustness against head motion

For the NeuroSynth and cross-species similarity analyses, we report the correlation of each contrast, with the corresponding difference in mean framewise displacement (Table S1 and Table S2). No such correlation is significant, for the cognitive matching results. However, for the human-macaque similarity results between sevoflurane anaesthesia and recovery, we do observe significant correlations with motion (Table S2). Nevertheless, correlations with motion are not observed when comparing anaesthesia against pre-anaesthesia wakefulness. For the fingerprinting analysis, we demonstrate that the results are not merely driven by the presence of high-motion participants. To do so, we repeat the analysis after applying a stringent criterion, excluding any participants with mean framewise displacement *<* 0.3 in any condition, resulting in the exclusion of N=3 participants, leaving N=12 for analysis. We show that the results are essentially unaltered, with anaesthesia significantly reducing both self-self similarity and differential identifiability (Fig. S13).

## DISCUSSION

Here we used pharmacological MRI under the effects of sevoflurane and propofol to determine whether anaesthetic-induced unconsciousness diminishes the uniqueness of the human brain: both with respect to the brains of other individuals, and the brain of another species entirely. We found that under deep anaesthesia, individual brains become less self-similar and less distinguishable from each other, in terms of functional connectivity. This effect is driven by reduced identifiability in transmodal association cortices.

These results are consistent with the notion that transmodal cortices, such as the default network and fronto-parietal control network, are particularly susceptible to anaesthesia, and loss of consciousness more generally [13, 46, 82]. In addition, association cortices exhibit the greatest rate of inter-individual variability [62]. This variability is not mere noise, however, since the frontoparietal and default networks consistently provide the largest contribution to identifiability in the awake resting brain [2, 88], indicating that their variability is individual-specific. This may be attributed to the fact that transmodal association cortices have the longest maturation times in the human brain, and the highest levels of synaptic plasticity and turnover [25, 49, 83]. Additionally, they also exhibit the lowest levels of intracortical myelination [17, 83], which is known to suppress plasticity both mechanically and chemically [26, 49]. As a result, transmodal cortices are relatively unconstrained by the underlying patterns of microstructure and anatomical connectivity [8, 74, 83, 91], and thus poised to change and adapt in response to environmental demands during the lifetime of each individual, which would account for their ability to encode individual-specific information in their connectivity. This is then temporarily (and reversibly) suppressed by anaesthesia, as the present results indicate.

We speculate that anaesthetic-induced suppression of individual differences in functional connectivity may be due to the consciousness-suppressing effects of anaesthesia. The default network in particular is well known to engage in both reflections about one’s own past and future, which by definition are unique to each individual [14, 15, 79]. By suppressing the idiosyncratic patterns of spontaneous thought that characterise the human brain even at rest, the present work indicates that anaesthetic-induced unconsciousness diminishes how such patterns are encoded in the macroscale activity and connectivity of the brain. Indeed, we found that as anaesthesia deepens, spontaneous brain activity is increasingly less well-characterised in terms of meta-analytic patterns pertaining to cognitive operations – whether automatically-defined or expert-curated. This effect is reversed upon recovery, despite the lingering presence of anaesthetic in the bloodstream.

Intriguingly, transmodal association cortices are not only the most heterogeneous between individuals, but also between species, exhibiting the greatest evolutionary expansion and the greatest expression of brain-related human-accelerated genes [16, 83, 92, 94]. We find that regional contributions to the anaesthetic-induced loss of identifiability are spatially correlated with both evolutionary cortical expansion and regional mean expression of human-accelerated genes. Furthermore, we showed that the anaesthetised human brain becomes more similar to the macaque brain in terms of regional functional connectivity an effect that is more evident at greater anaesthetic depth, and reversed upon recovery.

More broadly, the present results of diminished deviation between human and non-human primate un-der anaesthesia, are in line with previous work showing diminished deviation between structure and function under anaesthesia. Previous work had shown that across species, the anaesthetised brain’s patterns of timevarying functional connectivity become more similar to its underlying structural connectivity [7, 22, 32]. This phenomenon reflects diminished ability of the unconscious brain to engage in unusual patterns of connectivity that go beyond the dictates of anatomy. Intriguingly, psychedelics such as LSD and psilocybin (which induce hallucinations and highly bizarre subjective experiences) were recently found to have the opposite effect on structure-function coupling, making brain activity and connectivity less constrained by the underlying structural connectome [4, 45, 51]. Consistent with anaesthesia and psychedelics having opposite effects of structure-function relationships, a recent report suggests that psilocybin increases the idiosyncrasy of functional connectivity, resulting in greater differential identifiability [87] the opposite of what we found here with different anaesthetics. The present results suggest that the anaesthetised human brain is not only more similar to its underlying structure, but also more similar to the brains of other primates, with specifically human-expanded regions being the most affected by anaesthesia. Future research may investigate whether psychedelics have the opposite effect on the human brain, reducing in even greater difference between human and macaque especially since their primary molecular target, the 5HT2A receptor, is particularly prevalent in evolutionarily-expanded transmodal cortices [9].

This study also has a number of limitations that should be borne in mind. First, the small sample size, due to the technical and ethical challenges of performing anaesthesia in the scanner. While we replicated our results in a separate dataset, there is need in the field for larger sample sizes. Moreover, and encouragingly, identifiability and self-self similarity are higher between baseline wakefulness and recovery, which are the two scans most far apart in time—and everything else being equal, greater intervening time between scans would be expected to diminish identifiability. Additionally, the sevolfurane dataset was entirely comprised of male participants, and the majority of participants in the propofol dataset were also male: we look forward to future replications in sex-balanced datasets as the field expands. Second, we also acknowledge that the mapping of functional activation to psychological terms in NeuroSynth does not distinguish activations from deactivations [96]. However, we believe that our replication with meta-analytic maps defined using BrainMap [27] provides reassurance about the validity of our approach. Nonetheless, we note that the effects of cognitive matching from NeuroSynth - which are clearly dependent on depth of anaesthesia (Fig. 3) - were only observed for the sevoflurane dataset, which reaches deeper levels of anaesthesia. It will therefore be of particular interest to determine whether this result can be replicated in other datasets. In particular, this approach may prove valuable in datasets of patients with disorders of consciousness, where decoding brain responsiveness to task commands (e.g., “Imagine playing tennis”) has already enabled the identification of covert consciousness in behaviourally unresponsive patients [5, 19–21, 40, 61, 67, 68]. However, this paradigm requires patients’ ability to understand commands, keep them in working memory, and perform them - a non-trivial requirement for individuals who have suffered severe brain damage. To reduce this burden, researchers have also begun using spontaneous brain response to engaging narratives (e.g., clips from the movie “Taken”) [64]. However, this apparoach still requires language comprehension and working memory to follow the events. Decoding based on the match between meta-analytic maps and spontaenous brain activity without stimuli may further advance this line of research.

Altogether, the present results indicate that regardless of the specific anaesthetic used, anaesthetised human brains are less unique, both across individuals and even across species, with regions that are most heterogeneous across individuals and across species being especially affected.

## METHODS

### Sevoflurane data

The sevoflurane data included here have been published before [30, 47, 75], and we refer the reader to the original publication for details [75]. The ethics committee of the medical school of the Technische Universitat Munchen (Munchen, Germany) approved the current study, which was conducted in accordance with the Declaration of Helsinki. Written informed consent was obtained from volunteers at least 48 h before the study session. Twenty healthy adult men (20 to 36 years of age; mean, 26 years) were recruited through campus notices and personal contact, and compensated for their participation in the study. Before inclusion in the study, detailed information was provided about the protocol and risks, and medical history was reviewed to assess any previous neurologic or psychiatric disorder. A focused physical examination was performed, and a resting electrocardiogram was recorded. Further exclusion criteria were the following: physical status other than American Society of Anesthesiologists physical status I, chronic intake of medication or drugs, hardness of hearing or deafness, absence of fluency in German, known or suspected disposition to malignant hyperthermia, acute hepatic porphyria, history of halothane hepatitis, obesity with a body mass index more than 30 kg/m2, gastrointestinal disorders with a disposition for gastroesophageal regurgitation, known or suspected difficult airway, and presence of metal implants. Data acquisition took place between June and December 2013.

Sevoflurane concentrations were chosen so that participants tolerated artificial ventilation (reached at 2.0 vol%) and that burst-suppression (BS) was reached in all participants (around 4.4 vol%). To make group comparisons feasible, an intermediate concentration of 3.0 vol% was also used. In the MRI scanner, participants were in a resting state with eyes closed for 700s. Since EEG data were simultaneously acquired during MRI scanning [75] (though they are not analysed in the present study), visual online inspection of the EEG was used to verify that participants did not fall asleep during the pre-anaesthesia baseline scan. Sevoflurane mixed with oxygen was administered via a tight-fitting facemask using an fMRI-compatible anaesthesia machine (Fabius Tiro, Drager, Germany). Standard American Society of Anesthesiologists monitoring was performed: concentrations of sevoflurane, oxygen and carbon dioxide, were monitored using a cardiorespiratory monitor (DatexaS, General electric, USA). After administering an end-tidal sevoflurane concentration (etSev) of 0.4 vol% for 5 min, sevoflurane concentration was increased in a stepwise fashion by 0.2 vol% every 3 min until the participant became unconscious, as judged by the loss of responsiveness (LOR) to the repeatedly spoken command “squeeze my hand” two consecutive times. Sevoflurane concentration was then increased to reach an end-tidal concentration of approximately 3 vol%. When clinically indicated, ventilation was managed by the physician and a laryngeal mask suitable for fMRI (I-gel, Intersurgical, United Kingdom) was inserted. The fraction of inspired oxygen was then set at 0.8, and mechanical ventilation was adjusted to maintain end-tidal carbon dioxide at steady concentrations of 33±1.71 mmHg during BS, 34±1.12 mmHg during 3 vol%, and 33±1.49 mmHg during 2 vol% (throughout this article, mean±SD). Norepinephrine was given by continuous infusion (0.1±0.01*μg* kg-1 min1) through an intravenous catheter in a vein on the dorsum of the hand, to maintain the mean arterial blood pressure close to baseline values (baseline, 96±9.36 mmHg; BS, 88± 7.55 mmHg; 3 vol%, 88±8.4 mmHg; 2 vol%, 89±9.37 mmHg; follow-up, 98±9.41 mmHg). After insertion of the laryngeal mask airway, sevoflurane concentration was gradually increased until the EEG showed burst-suppression with suppression periods of at least 1,000 ms and about 50% suppression of electrical activity (reached at 4.34±0.22 vol%), which is characteristic of deep anaesthesia. At that point, another 700s of electroencephalogram and fMRI was recorded. Further 700s of data were acquired at steady end-tidal sevoflurane concentrations of 3 and 2 vol%, respectively (corresponding to Ramsay scale level 6), each after an equilibration time of 15 min. In a final step, etSev was reduced to two times the concentration at LOR. However, most of the participants moved or did not tolerate the laryngeal mask any more under this condition: therefore, this stage was not included in the analysis. Sevoflurane administration was then terminated, and the scanner table was slid out of the MRI scanner to monitor post-anaesthetic recovery. The participants was manually ventilated until spontaneous ventilation returned. The laryngeal mask was removed as soon as the patient opened his mouth on command. The physician regularly asked the participant to squeeze their hand: recovery of responsiveness was noted to occur as soon as the command was followed. Fifteen minutes after the time of recovery of responsiveness, the Brice interview was administered to assess for awareness during sevoflurane exposure; the interview was repeated on the phone the next day. After a total of 45 min of recovery time, another resting-state combined fMRI-EEG scan was acquired (with eyes closed, as for the baseline scan). When participants were alert, oriented, cooperative, and physiologically stable, they were taken home by a family member or a friend appointed in advance.

Although the original study acquired both functional MRI (fMRI) and electroencephalographic (EEG) data, in the present work we only considered the fMRI data. Data acquisition was carried out on a 3-Tesla magnetic resonance imaging scanner (Achieva Quasar Dual 3.0T 16CH, The Netherlands) with an eight-channel, phased-array head coil. The data were collected using a gradient echo planar imaging sequence (echo time = 30 ms, repetition time (TR) = 1.838 s, flip angle = 75 deg, field of view = 220×220 mm2, matrix = 72×72, 32 slices, slice thickness = 3 mm, and 1 mm interslice gap; 700-s acquisition time, resulting in 350 functional volumes). The anatomical scan was acquired before the functional scan using a T1-weighted MPRAGE sequence with 240× 240 ×170 voxels (1×1×1 mm voxel size) covering the whole brain. A total of 16 volunteers completed the full protocol and were included in our analyses; one participant was excluded due to high motion, leaving N=15 for analysis.

### Propofol data

The propofol data were collected between May and November 2014 at the Robarts Research Institute, Western University, London, Ontario (Canada), and have been published before [46, 65]. The study received ethical approval from the Health Sciences Research Ethics Board and Psychology Research Ethics Board of Western University (Ontario, Canada). Healthy volunteers (n=19) were recruited (18–40 years; 13 males). Volunteers were right-handed, native English speakers, and had no history of neurological disorders. In accordance with relevant ethical guidelines, each volunteer provided written informed consent, and received monetary compensation for their time. Due to equipment malfunction or physiological impediments to anaesthesia in the scanner, data from n=3 participants (1 male) were excluded from analyses, leaving a total n=16 for analysis.

Resting-state fMRI data were acquired at different propofol levels: no sedation (Awake), Deep anaesthesia (corresponding to Ramsay score of 5) and also dur-ing post-anaesthetic recovery. As previously reported [46], for each condition fMRI acquisition began after two anaesthesiologists and one anaesthesia nurse independently assessed Ramsay level in the scanning room. The anaesthesiologists and the anaesthesia nurse could not be blinded to experimental condition, since part of their role involved determining the participants’ level of anaesthesia. Note that the Ramsay score is designed for critical care patients, and therefore participants did not receive a score during the Awake condition before propofol administration: rather, they were required to be fully awake, alert and communicating appropriately. To provide a further, independent evaluation of participants’ level of responsiveness, they were asked to perform two tasks: a test of verbal memory recall, and a computerbased auditory target-detection task. Wakefulness was also monitored using an infrared camera placed inside the scanner. Propofol was administered intravenously using an AS50 auto syringe infusion pump (Baxter Healthcare, Singapore); an effect-site/plasma steering algorithm combined with the computer-controlled infusion pump was used to achieve step-wise sedation increments, followed by manual adjustments as required to reach the desired target concentrations of propofol according to the TIVA Trainer (European Society for Intravenous Aneaesthesia, eurosiva.eu) pharmacokinetic simulation program. This software also specified the blood concentrations of propofol, following the Marsh 3compartment model, which were used as targets for the pharmacokinetic model providing target-controlled infusion. After an initial propofol target effect-site concentration of 0.6 *μ*g mL-1, concentration was gradually increased by increments of 0.3 *μ*g mL1, and Ramsay score was assessed after each increment: a further increment occurred if the Ramsay score was lower than 5. The mean estimated effect-site and plasma propofol concentrations were kept stable by the pharmacokinetic model delivered via the TIVA Trainer infusion pump. Ramsay level 5 was achieved when participants stopped responding to verbal commands, were unable to engage in conversation, and were rousable only to physical stimulation. Once both anaesthesiologists and the anaesthesia nurse all agreed that Ramsay sedation level 5 had been reached, and participants stopped responding to both tasks, data acquisition was initiated. The mean estimated effect-site propofol concentration was 2.48 (1.82-3.14) *μ*g mL-1, and the mean estimated plasma propofol concentration was 2.68 (1.92-3.44) *μ*g mL-1. Mean total mass of propofol administered was 486.58 (373.30-599.86) mg. These values of variability are typical for the pharmacokinetics and pharmacodynamics of propofol.

Oxygen was titrated to maintain SpO2 above 96%. At Ramsay 5 level, participants remained capable of spontaneous cardiovascular function and ventilation. However, the sedation procedure did not take place in a hospital setting; therefore, intubation during scanning could not be used to ensure airway security during scanning. Consequently, although two anaesthesiologists closely monitored each participant, scanner time was minimised to ensure return to normal breathing following deep sedation. No state changes or movement were noted during the deep sedation scanning for any of the participants included in the study. Propofol was discontinued following the deep anaesthesia scan, and participants reached level 2 of the Ramsey scale approximately 11 minutes afterwards, as indicated by clear and rapid responses to verbal commands. This corresponds to the “recovery” period. As previously reported [46], once in the scanner participants were instructed to relax with closed eyes, without falling asleep. Resting-state functional MRI in the absence of any tasks was acquired for 8 minutes for each participant. A further scan was also acquired during auditory presentation of a plot-driven story through headphones (5-minute long). Participants were instructed to listen while keeping their eyes closed. The present analysis focuses on the resting-state data only; the story scan data have been published separately [65] and will not be discussed further here.

As previously reported [46], MRI scanning was performed using a 3-Tesla Siemens Tim Trio scanner (32channel coil), and 256 functional volumes (echo-planar images, EPI) were collected from each participant, with the following parameters: slices = 33, with 25% interslice gap; resolution = 3mm isotropic; TR = 2000ms; TE= 30ms; flip angle = 75 degrees; matrix size = 64×64. The order of acquisition was interleaved, bottomup. Anatomical scanning was also performed, acquiring a high-resolution T1weighted volume (32-channel coil, 1mm isotropic voxel size) with a 3D MPRAGE sequence, using the following parameters: TA = 5min, TE = 4.25ms, 240×256 matrix size, 9 degrees flip angle [46].

### FMRI preprocessing and denoising

We applied a standard preprocessing pipeline in accordance with our previous publications with anaesthesia data [46, 47]. Preprocessing was performed using the *CONN* toolbox, version 17f (CONN; http://www.nitrc.org/projects/conn) [93], implemented in MATLAB 2016a. The pipeline involved the following steps: removal of the first 10s, to achieve steady-state magnetization; motion correction; slice-timing correction; identification of outlier volumes for subsequent scrubbing by means of the quality assurance/artifact rejection software *art* (http://www.nitrc.org/projects/artifact_detect); normalisation to Montreal Neurological Institute (MNI-152) standard space (2 mm isotropic resampling resolution), using the segmented grey matter image from each participant’s T1-weighted anatomical image, together with an a priori grey matter template.

Denoising was also performed using the CONN toolbox, using the same approach as in our previous publications with pharmaco-MRI datasets [46, 47]. Pharmacological agents can induce alterations in physiological parameters (heart rate, breathing rate, motion) or neurovascular coupling. The anatomical CompCor (aCompCor) method removes physiological fluctuations by extracting principal components from regions unlikely to be modulated by neural activity; these components are then included as nuisance regressors [95]. Following this approach, five principal components were extracted from white matter and cerebrospinal fluid signals (using individual tissue masks obtained from the T1-weighted structural MRI images) [93]; and regressed out from the functional data together with six individual-specific realignment parameters (three translations and three rotations) as well as their first-order temporal derivatives; followed by scrubbing of outliers identified by ART, using Ordinary Least Squares regression [93]. Finally, the denoised BOLD signal timeseries were linearly detrended and band-pass filtered to eliminate both low-frequency drift effects and high-frequency noise, thus retaining frequencies between 0.008 and 0.09 Hz. The step of global signal regression (GSR) has received substantial attention in the literature as a denoising method [63, 72, 73]. However, recent work has demonstrated that the global signal contains behaviourally relevant information [44] and, crucially, information about states of consciousness, across pharmacological and pathological perturbations [84]. Therefore, in line with ours and others’ previous studies, here we avoided GSR in favour of the aCompCor denoising procedure, which is among those recommended.

Finally, denoised BOLD signals were parcellated into 200 cortical regions from the Schaefer atlas [76]. We also replicated our results with the anatomical Desikan-Killiany cortical parcellation [23], and with a combined cortical-subcortical atlas comprising 200 cortical ROIs from the Schaefer atlas, and additional 32 subcortical ROIs from the subcortical atlas of Tian and colleagues [86], as previously recommended [50]. For comparison with the macaque data, a human-adapted version of the 82-ROI cortical parcellation of Kotter and Wanke was used [41], as adapted by [12]. Functional connectivity was estimated for each individual and each condition, as the Pearson correlation between between pairs of denoised and parcellated BOLD timeseries.

### Macaque fMRI data

The non-human primate MRI data were made available as part of the Primate neuroimaging Data-Exchange (PRIME-DE) monkey MRI data sharing initiative, a recently introduced open resource for non-human primate imaging [57].

The data preprocessing and denoising followed the same procedures as in our previous publication [48]. We used fMRI data from rhesus macaques (*Macaca mulatta*) scanned at Newcastle University. This samples includes 14 exemplars (12 male, 2 female); Age distribution: 3.9-13.14 years; Weight distribution: 7.2-18 kg (full sample description available online: http://fcon_1000.projects.nitrc.org/indi/PRIME/files/newcastle.csv and http://fcon_1000.projects.nitrc.org/indi/PRIME/newcastle.html).

Ethics approval: All of the animal procedures performed were approved by the UK Home Office and comply with the Animal Scientific Procedures Act (1986) on the care and use of animals in research and with the European Directive on the protection of animals used in research (2010/63/EU). We support the Animal Research Reporting of In Vivo Experiments (ARRIVE) principles on reporting animal research. All persons involved in this project were Home Office certified and the work was strictly regulated by the U.K. Home Office. Local Animal Welfare Review Body (AWERB) approval was obtained. The 3Rs principles compliance and assessment was conducted by National Centre for 3Rs (NC3Rs). Animal in Sciences Committee (UK) approval was obtained as part of the Home Office Project License approval. Animal care and housing: All animals were housed and cared for in a group-housed colony, and animals performed behavioural training on various tasks for auditory and visual neuroscience. No training took place prior to MRI scanning. Macaque MRI acquisition

Animals were scanned in a vertical Bruker 4.7T primate dedicated scanner, with single channel or 4-8 channel parallel imaging coils used. No contrast agent was used. Optimization of the magnetic field prior to data acquisition was performed by means of 2nd order shim, Bruker and custom scanning sequence optimization. Animals were scanned upright, with MRI compatible headpost or non-invasive head immobilisation, and working on tasks or at rest (here, only resting-state scans were included). Eye tracking, video and audio monitoring were employed during scanning. Resting-state scanning was performed for 21.6 minutes, with a TR of 2600ms, 17ms TE, Effective Echo Spacing of 0.63ms, voxels size 1.22× 1.22×1.24. Phase Encoding Direction: Encoded in columns. Structural scans comprised a T1 structural, MDEFT sequence with the following parameters: TE: 6ms; TR: 750 ms; Inversion delay: 700ms; Number of slices: 22; In-plane field of view: 12.8×9.6 cm^2^ on a grid of 256×192 voxels; Voxel resolution: 0.5×0.5×2mm; Number of segments: 8.

The macaque MRI data were preprocessed using the recently developed pipeline for non-human primate MRI analysis, Pypreclin, which addresses several specificities of monkey research. The pipeline is described in detail in the associated publication [85]. Briefly, it includes the following steps: (i) Slice-timing correction. (ii) Correction for the motion-induced, time-dependent B0 inhomogeneities. (iii) Reorientation from acquisition position to template; here, we used the recently developed National Institute of Mental Health Macaque Template (NMT): a high-resolution template of the average macaque brain generated from in vivo MRI of 31 rhesus macaques (Macaca mulatta) [37]. (iv) Realignment to the middle volume using FSL MCFLIRT function. (v) Normalisation and masking using Joe’s Image Program (JIP-align) routine (http://www.nmr.mgh.harvard.edu/~jbm/jip/, Joe Mandeville, Massachusetts General Hospital, Harvard University, MA, USA), which is specifically designed for preclinical studies: the normalization step aligns (affine) and warps (non-linear alignment using distortion field) the anatomical data into a generic template space. (vi) B1 field correction for low-frequency intensity non-uniformities present in the data. (vii) Coregistration of functional and anatomical images, using JIP-align to register the mean functional image (moving image) to the anatomical image (fixed image) by applying a rigid transformation. The anatomical brain mask was obtained by warping the template brain mask using the deformation field previously computed during the normalization step. Then, the functional images were aligned with the template space by composing the normalization and coregistration spatial transformations.

#### Denoising

The aCompCor denoising method implemented in the CONN toolbox was used to denoise the macaque functional MRI data, to ensure consistency with the human data analysis pipeline. White matter and CSF masks were obtained from the corresponding probabilistic tissue maps of the high-resolution NMT template (eroded by 1 voxel); their first five principal components were regressed out of the functional data, as well as linear trends and 6 motion parameters (3 translations and 3 rotations) and their first derivatives. To make human and macaque data comparable, the macaque data were also bandpass filtered in the same 0.008-0.09 Hz range used for human data.

Out of the 14 total animals present in the Newcastle sample, 10 had available awake resting-state fMRI data; of these 10, all except the first animal had two scanning sessions available. Thus, the total was 19 distinct sessions across 10 individual macaques. We combined them to obtain a map of regional macaque FC.

### Brain fingerprinting

“Brain fingerprinting” refers to using brain-derived metrics (here: the functional connectivity obtained from resting-state functional MRI) to discriminate individuals from each other, analogously to how the grooves on one’s fingertips may be used to discern one’s identity. This requires brain fingerprints (just like conventional fingerprints) to be different across different people (to avoid confusing distinct individuals) but consistent within the same individual (to track identity).

Let A be the “identifiability matrix”, i.e. the (square, non symmetric) matrix of similarity between individuals’ *test* and *retest* scans, such that the size of A is S-by-S (with S being the number of individuals in the dataset). Each entry of A is obtained as the correlation between the corresponding individuals’ vectorised matrices of parcellated functional connectivity. Let *I*_self_ = ⟨*a*_*ii*_⟩represent the average of the main diagonal elements of A, which consist of the Pearson correlation values between scans of same individual: from now on, we will refer to this quantity as self-identifiability or *I*_self_. Similarly, let *I*_others_ = ⟨*a*_*ij*_ ⟩define the average of the off-diagonal elements of matrix A, i.e. the correlation between scans of different individuals *i* and *j*. Then, we define the differential identifiability (*I*_diff_) of the sample as the difference between both terms:

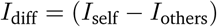

(with *i* ≠ *j*) which quantifies the difference between the average within-participant FCs similarity and the average between-participants FCs similarity. The higher the value of *I*_diff_, the higher the individual fingerprint overall along the population [2].

We can also quantify the edgewise identifiability of individuals by using intra-class correlation (ICC; [2]). ICC is a widely used measure in statistics, most commonly to assess the percent of agreement between units (or ratings/scores) of different groups (or raters/judges) [6, 56, 78]. It describes how strongly units in the same group resemble each other. The stronger the agreement between the ratings, the higher its ICC value. We use ICC to quantify to which extent the connectivity value of an edge (functional connectivity value between two brain regions) could separate within and between participants. In other words, the higher the ICC, the higher the identifiability of the connectivity edge [2].

### Meta-analytic cognitive matching from NeuroSynth

Continuous measures of the association between voxels and cognitive categories were obtained from NeuroSynth, an automated term-based meta-analytic tool that synthesizes results from more than 14 000 published fMRI studies by searching for high-frequency key words (such as “pain” and “attention” terms) that are systematically mentioned in the papers alongside fMRI voxel coordinates (https://github.com/neurosynth/neurosynth), using the volumetric association test maps [96]. This measure of association strength is the tendency that a given term is reported in the functional neuroimaging study if there is activation observed at a given voxel. Note that NeuroSynth does not distinguish between areas that are activated or deactivated in relation to the term of interest, nor the degree of activation, only that certain brain areas are frequently reported in conjunction with certain words. Although more than a thousand terms are catalogued in the NeuroSynth engine, we refine our analysis by focusing on cognitive function and therefore we limit the terms of interest to cognitive and behavioural terms. To avoid introducing a selection bias, we opted for selecting terms in a data-driven fashion instead of selecting terms manually. Therefore, terms were selected from the Cognitive Atlas, a public ontology of cognitive science [69], which includes a comprehensive list of neurocognitive terms. This approach totaled to *t* = 123 terms, ranging from umbrella terms (“attention”, “emotion”) to specific cognitive processes (“visual attention”, “episodic memory”), behaviours (“eating”, “sleep”), and emotional states (“fear”, “anxiety”) (note that the 123 term-based meta-analytic maps from NeuroSynth do not explicitly exclude patient studies). The Cognitive Atlas subdivision has previously been used in conjunction with NeuroSynth [1, 33, 53], so we opted for the same approach to make our results comparable to previous reports. The probabilistic measure reported by NeuroSynth can be interpreted as a quantitative representation of how regional fluctuations in activity are related to psychological processes. As with the restingstate BOLD data, voxelwise NeuroSynth maps were parcellated into 200 cortical regions according to the Schaefer atlas [76] (or 68 for the replication with Desikan-Killiany atlas, or 232 cortical and subcortical regions for replication with subcortex included).

For each individual, their parcellated BOLD signals at each point in time were spatially correlated against each NeuroSynth map, producing one value of correlation per NeuroSynth map, per BOLD volume. We refer to this operation as “cognitive matching”. For each volume, the quality of cognitive matching was quantified as the highest value of (positive) correlation across all maps. These values were subsequently averaged across all volumes to obtain a single value per condition, per participant. As an alternative, instead of using the highest positive correlation, we also considered the mean magnitude of correlation (regardless of sign) across *all* maps, subsequently averaging across volumes as described above.

We also repeated the cognitive matching separately for NeuroSynth maps exhibiting a positive spatial correlation with the “archetypal axis” of [83] (see below), i.e. those primarily exhibiting positive values in transmodal association cortex; and for maps exhibiting a negative correlation with the archetypal axis, which primarily comprised activation in unimodal (sensory) cortices.

### Alternative meta-analytic matching from BrainMap

Whereas NeuroSynth is an automated tool, BrainMap is an expert-curated repository: it includes the brain coordinates that are significantly activated during thousands of different experiments from published neuroimaging studies [27, 43]. As a result, NeuroSynth terms and BrainMap behavioural domains differ considerably. Here, we used maps in the Desikan-Killiany atlas, pertaining to 66 unique behavioural domains (the same as in [33]), obtained from 8, 703 experiments. Experiments conducted on unhealthy participants were excluded, as well as experiments without a defined behavioural domain.

### Archetypal axis, cortical expansion, inter-individual variability, and human-accelerated gene expression maps

To contextualise our regional pattern of anaesthetic-induced changes in identifiability, we obtained relevant brain maps from the literature using *neuromaps* (https://netneurolab.github.io/neuromaps/). We fetched and parcellated the map of sensory-association archetypal axis from [83], the map of cortical expansion between macaque and human from [94], and the map of interindividual variability of functional connectivity from [62].

Human-accelerated genes are genes associated with so-called human accelerated regions of the human genome, identified as set of loci that displayed accelerated divergence in the human lineage by comparing the human genome with that of the chimpanzee (*Pan troglodytes*), one of our closest living evolutionary relatives [70, 71]. Among these human-accelerated genes,”HAR-brain genes” pertain to brain function and development [92]. The map of mean regional expression of HAR-brain gene expression was obtained as follows. First, the list of 415 HAR-brain genes was obtained from [92] (see the original publication for details of how these genes were selected). Then, regional gene expression for each of the 200 cortical regions of the Schaefer atlas was obtained using the *abagen* toolbox https://abagen.readthedocs.io/ [52], following *abagen*’s default processing workflow, and mirroring data between homologous cortical regions to ensure adequate coverage of both left (data from six donors) and right hemisphere (data from two donors). Distances between samples were evaluated on the cortical surface with a 2 mm distance threshold. Gene expression data were normalised across the cortex using outlier-robust sigmoid normalisation. Out of the resulting 15, 633 genes, 392 were among the list of HAR-brain genes from Wei and colleagues. Finally, the map of regional mean expression of HAR-brain genes was obtained as the regional mean normalised gene expression across the 392 genes.

### Human structural connectome from Human Connectome Project

We used diffusion MRI (dMRI) data from the 100 unrelated participants (54 females and 46 males, mean age = 29.1±3.7 years) of the HCP 900 participants data release [89]. All HCP scanning protocols were approved by the local Institutional Review Board at Washington University in St. Louis. The diffusion weighted imaging (DWI) acquisition protocol is covered in detail elsewhere [28]. The diffusion MRI scan was conducted on a Siemens 3T Skyra scanner using a 2D spin-echo singleshot multiband EPI sequence with a multi-band factor of 3 and monopolar gradient pulse. The spatial resolution was 1.25 mm isotropic. TR=5500 ms, TE=89.50ms. The b-values were 1000, 2000, and 3000 s/mm^2^. The total number of diffusion sampling directions was 90, 90, and 90 for each of the shells in addition to 6 b0 images. We used the version of the data made available in DSI Studio-compatible format at http://brain.labsolver.org/ diffusion-mri-templates/hcp-842-hcp-1021 [97].

We adopted previously reported procedures to reconstruct the human connectome from DWI data. The minimally-preprocessed DWI HCP data [28] were corrected for eddy current and susceptibility artifact. DWI data were then reconstructed using q-space diffeomorphic reconstruction (QSDR [99]), as implemented in DSI Studio (www.dsi-studio.labsolver.org). QSDR calculates the orientational distribution of the density of diffusing water in a standard space, to conserve the diffusible spins and preserve the continuity of fiber geometry for fiber tracking. QSDR first reconstructs diffusion-weighted images in native space and computes the quantitative anisotropy (QA) in each voxel. These QA values are used to warp the brain to a template QA volume in Montreal Neurological Institute (MNI) space using a nonlinear registration algorithm implemented in the statistical parametric mapping (SPM) software. A diffusion sampling length ratio of 2.5 was used, and the output resolution was 1 mm. A modified FACT algorithm [98] was then used to perform deterministic fiber tracking on the reconstructed data, with the following parameters [50]: angular cutoff of 55 ^*?*^(Fig. **??**), step size of 1.0 mm, minimum length of 10 mm, maximum length of 400mm, spin density function smoothing of 0.0, and a QA threshold determined by DWI signal in the cerebrospinal fluid. Each of the streamlines generated was automatically screened for its termination location. A white matter mask was created by applying DSI Studio’s default anisotropy threshold (0.6 Otsu’s threshold) to the spin distribution function’s anisotropy values. The mask was used to eliminate streamlines with premature termination in the white matter region. Deterministic fiber tracking was performed until 1, 000, 000 streamlines were reconstructed for each individual.

For each individual, their structural connectome was reconstructed by drawing an edge between each pair of regions *i* and *j* from the Schaefer cortical atlas [76] if there were white matter tracts connecting the corresponding brain regions end-to-end; edge weights were quantified as the number of streamlines connecting each pair of regions, normalised by ROI distance and size.

A group-consensus matrix *A* across participants was then obtained using the distance-dependent procedure of Betzel and colleagues, to mitigate concerns about inconsistencies in reconstruction of individual participants’ structural connectomes [11]. This approach seeks to preserve both the edge density and the prevalence and length distribution of inter- and intra-hemispheric edge length distribution of individual participants’ connectomes, and it is designed to produce a representative connectome [11, 59]. This procedure produces a binary consensus network indicating which edges to preserve. The final edge density was 27%. The weight of each nonzero edge is then computed as the mean of the corresponding non-zero edges across participants.

### Statistical analyses

The statistical significance of differences between conditions (here: levels of anaesthesia) was determined with permutation t-tests (paired-sample), with 10 000 permutations. The use of permutation tests alleviated the need to assume normality of data distributions (which was not formally tested). All tests were two-sided, with an *α* value of 0.05. The effect sizes were estimated using Hedge’s measure of the standardized mean difference, *g*, which is interpreted in the same way as Cohen’s *d*, but more appropriate for small sample sizes [34]. The Measures of Effect Size Toolbox for MATLAB https://github.com/hhentschke/measures-of-effect-size-toolbox was used [35]. Anaesthesia conditions were compared separately against wakefulness and against recovery, and the false positive rate against multiple comparisons was controlled using the false discovery rate (FDR) correction [10], separately for these two cases. The statistical significance of spatial correlation between brain maps was assessed non-parametrically via comparison against a null distribution of null maps with preserved spatial autocorrelation [53, 90].

## Data availability

The original pharmacological fMRI data are available from the corresponding authors of the original publications referenced herein. The Allen Human Brain Atlas transcriptomic database is available at https://human.brain-map.org/; NeuroSynth is available at https://neurosynth.org/. The list of human-accelerated brain genes is available from the Supplementary Material of [92]. The Newcastle macaque fMRI data are available from the PRIMEDE database (http://fcon_1000.projects.nitrc.org/indi/indiPRIME.html). Diffusion MRI data for the Human Connectome Project in DSI Studio-compatible format are available at http://brain.labsolver.org/diffusion-mri-templates/hcp-842-hcp-1021.

## Code availability

Code for brain fingerprinting is freely available at https://github.com/eamico/MEG_fingerprints. The code for spin-based permutation testing of cortical correlations is freely available at https://github.com/frantisekvasa/rotate_parcellation. *DSI Studio* is freely available at https://dsi-studio.labsolver.org/. The *CONN* toolbox is freely available at http://www.nitrc.org/projects/conn. The Pypreclin code is available at https://github.com/neurospin/pypreclin. The *abagen* toolbox is available at https://abagen.readthedocs.io/ The *neuromaps* toolbox is available at https://netneurolab.github.io/neuromaps/.

## Acknowledgments

We wish to express our gratitude to the PRIME-DE initiative, to the organizers and managers of PRIME-DE and to all the institutions that contributed to the PRIMEDE database (http://fcon_1000.projects.nitrc.org/indi/indiPRIME.html), with special thanks to the Newcastle team. We are also grateful to A. Grigis, J. Tasserie and B. Jarraya for their assistance with the Pypreclin code. We are also grateful to Dr Kelly Shen for sharing the human version of the RM macaque parcellation. We thank members of the Network Neuroscience Lab for helpful discussion.

## Author contributions

A.I.L., E.A. and B.M. conceived of the study idea. A.I.L. performed the formal analysis. D.G., R.I., A.R., D.J., L.N., A.M.O., E.A.S., D.B., contributed data. E.A.S., E.A. contributed to interpretation. A.I.L. and B.M. wrote the manuscript with inputs from all coauthors.

