## Supplementary Figures and Tables for "General anaesthesia reduces the uniqueness of brain connectivity across individuals and across species"

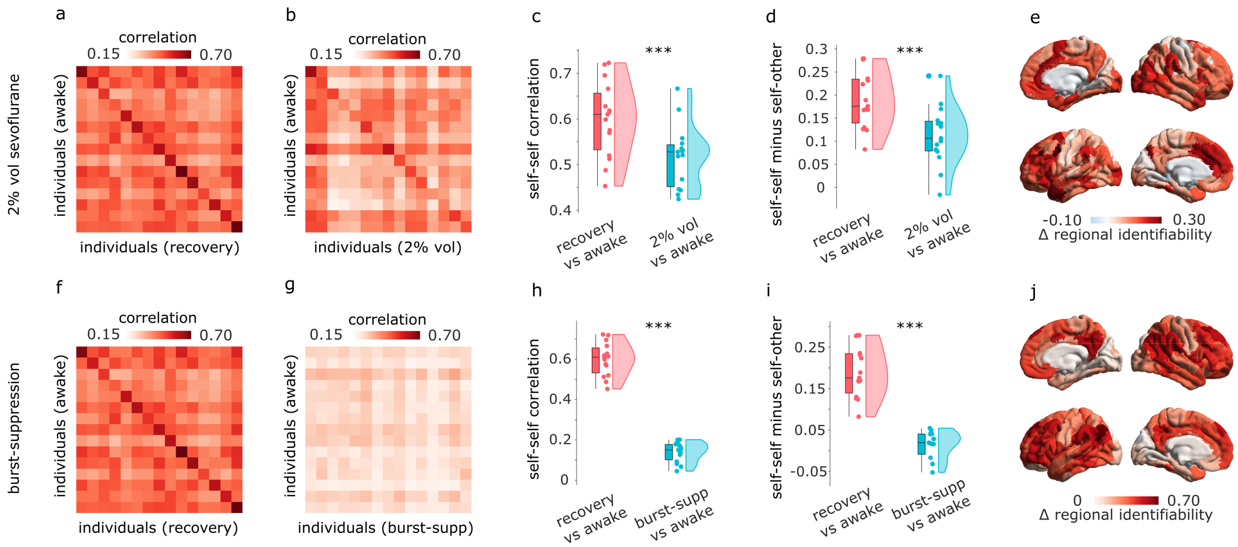


Figure S1. **Replication of identifiability results at different doses of sevoflurane |** (**a**) Identifiability matrix between wakefulness and post-anaesthetic recovery. (**b**) Identifiability matrix between wakefulness and vol 2% sevoflurane anaesthesia (right). Entries along the diagonal, represent self-self similarity (correlation of FC patterns), whereas off-diagonal entries represent self-other similarity. (**c**) Self-self similarity is significantly higher between two conscious states, than between wakefulness and vol 2% sevoflurane. (**d**) The difference between self-self correlation and mean self-other correlation (differential identifiability) is significantly higher between two conscious states, than between wakefulness and vol 2% sevoflurane. (**e**) The regional distribution of contributions to identifiability (change in intra-class correlation coefficient) is plotted on the cortical surface. It is significantly spatially correlated with the corresponding map obtained with vol 3% sevoflurane: Spearman $\rho = 0.61$, $p_{spin} < 0.001$, $N = 200$ regions. (**f**) Identifiability matrix between wakefulness and post-anaesthetic recovery. (**g**) Identifiability matrix between wakefulness and burst-suppression level of sevoflurane anaesthesia. (**h**) Self-self similarity is significantly higher between two conscious states, than between wakefulness and burst-suppression level of sevoflurane. (**i**) The difference between self-self correlation and mean self-other correlation (differential identifiability) is significantly higher between two conscious states, than between wakefulness and burst-suppression level of sevoflurane. (**j**) The regional distribution of contributions to identifiability (change in intra-class correlation coefficient) is plotted on the cortical surface. It is significantly spatially correlated with the corresponding map obtained with vol 3% sevoflurane: Spearman $\rho = 0.80$, $p_{spin} < 0.001$, $N = 200$ regions. ***, $p < 0.001$.

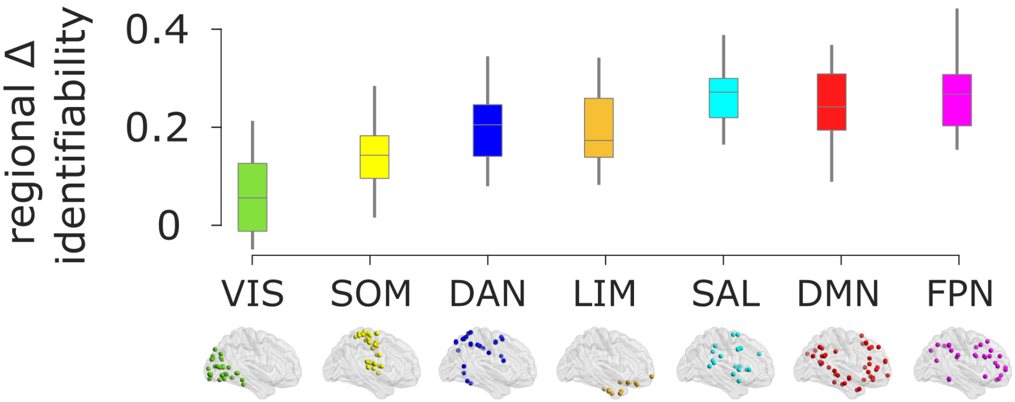


Figure S2. **Regional identifiability across intrinsic connectivity networks |**. SOM, somatomotor network; VIS, visual network; SAL, salience network; DAN, dorsal attention network; FPN, fronto-parietal control network; DMN, default mode network. Boxplots: center line, median; box limits, upper and lower quartiles; whiskers, $1.5\times$ interquartile range.

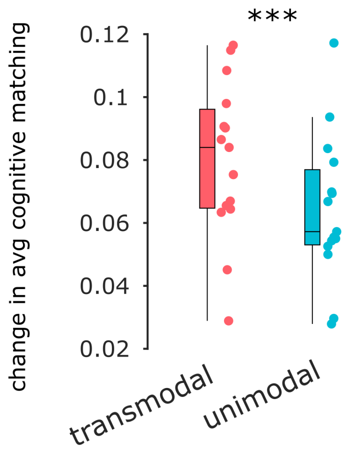


Figure S3. **Change in cognitive matching for unimodal versus transmodal ends of the cortical hierarchy|**. NeuroSynth maps were correlated with the archetypal axis map of [83]; maps exhibiting a positive correlation were included at the transmodal end, whereas maps correlating negatively were included at the unimodal end. Box-plot: center line, median; box limits, upper and lower quartiles; whiskers, $1.5\times$ interquartile range. ***, $p < 0.001$.

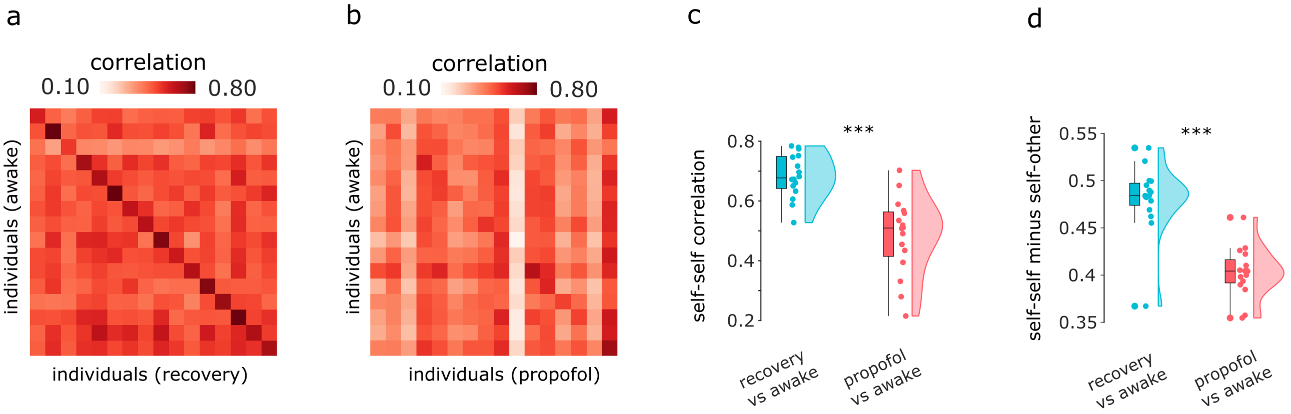


Figure S4. **Identifiability of individual connectomes is diminished under propofol anaesthesia.** (**a**) Identifiability matrix between wakefulness and post-anaesthetic recovery. (**b**) Identifiability matrix between wakefulness and propofol anaesthesia (right). Entries along the diagonal, represent self-self similarity (correlation of FC patterns), whereas off-diagonal entries represent self-other similarity. (**c**) Self-self similarity is significantly higher between two conscious states, than between wakefulness and propofol anaesthesia. (**d**) The difference between self-self correlation and mean self-other correlation (differential identifiability) is significantly higher between two conscious states, than between wakefulness and propofol anaesthesia. Box-plot: center line, median; box limits, upper and lower quartiles; whiskers, $1.5\times$ interquartile range. ***, $p < 0.001$.

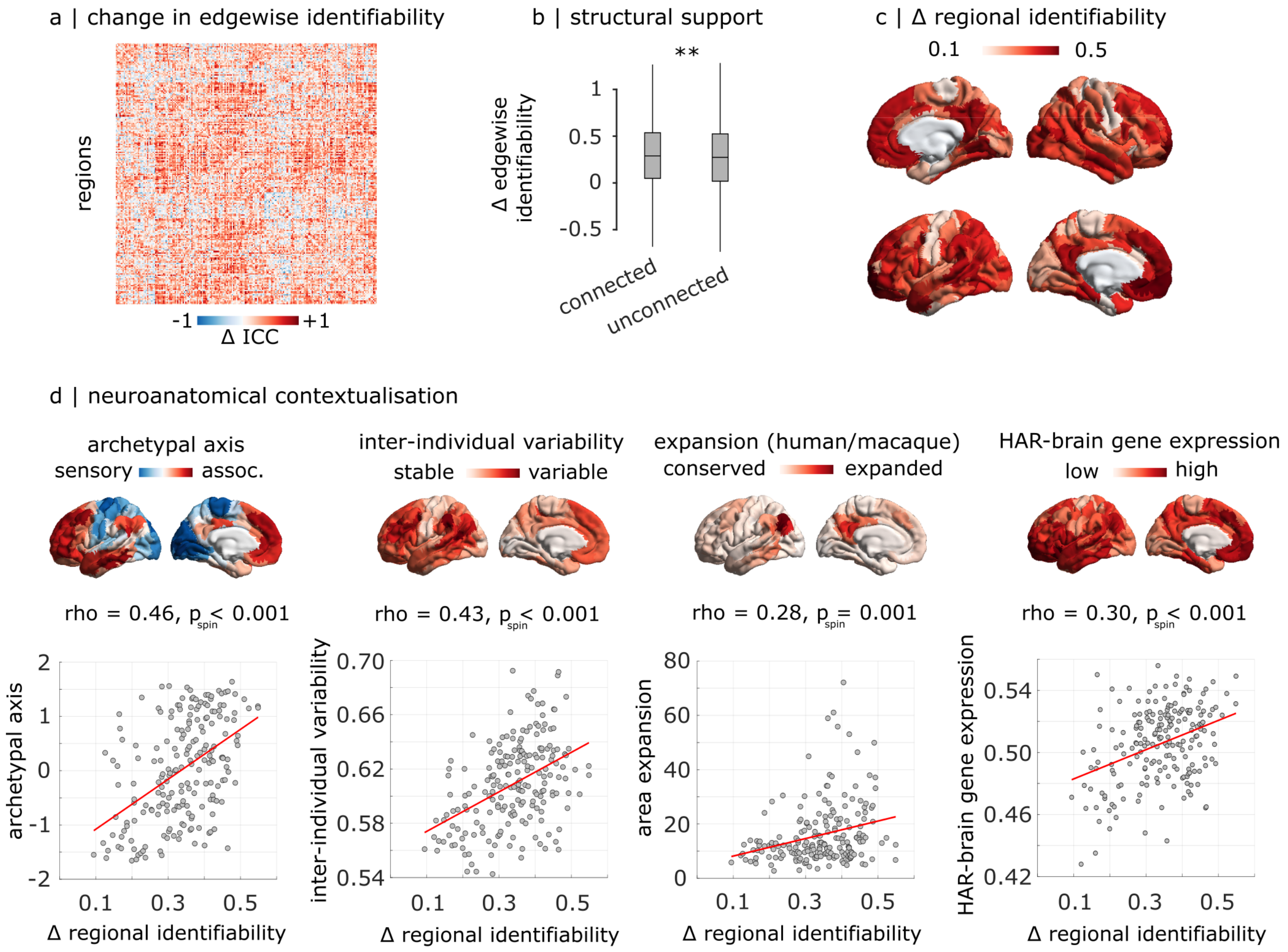


Figure S5. **Anatomical characterisation of contributions to propofol-induced loss of identifiability** | (**a**) Edge-level difference in intra-class correlation coefficient between awake-recovery and awake-propofol. (**b**) The anaesthetic-induced loss of ICC is significantly more pronounced for functional connections with an underlying structural direct connection, than without. Box-plot: center line, median; box limits, upper and lower quartiles; whiskers, $1.5\times$ interquartile range. **, $p < 0.01$. (**c**) Regional distribution of propofol-induced loss of ICC, projected onto the cortical surface. It is significantly spatially correlated with the corresponding map obtained with sevoflurane: Spearman $\rho = 0.35$, $p_{spin} < 0.001$, $N = 200$ regions. (**d**) The propofol-induced regional loss of ICC is significantly spatially aligned with the archetypal sensory-association axis of cortical organisation; the regional distribution of inter-individual variability of functional connectivity; the regional distribution of cortical expansion between macaque and human brains; and the regional expression of human-accelerated genes pertaining to brain function and development ("HAR-brain genes"); $N = 200$ regions

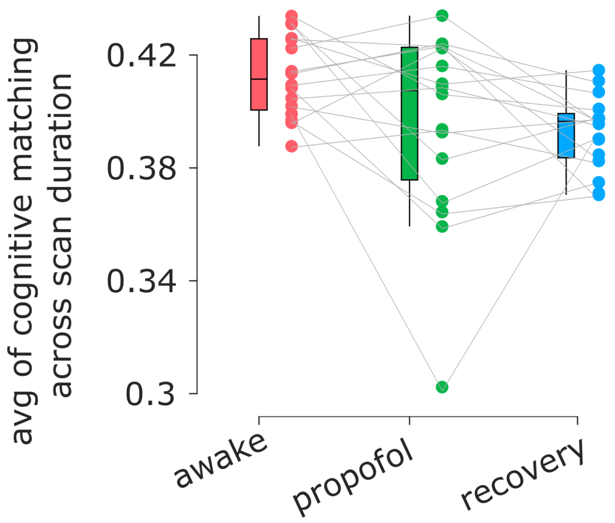


Figure S6. **Cognitive matching from brain activity under wakefulness and propofol anaesthesia |** Ordinate: mean across time of the best decoding score (maximum spatial correlation between brain activity and 123 NeuroSynth meta-analytic maps). *** (black), $p < 0.001$ against wakefulness (FDR-corrected); *** (gray), $p < 0.001$ against recovery (FDR-corrected). Box-plot: center line, median; box limits, upper and lower quartiles; whiskers, $1.5\times$ interquartile range.

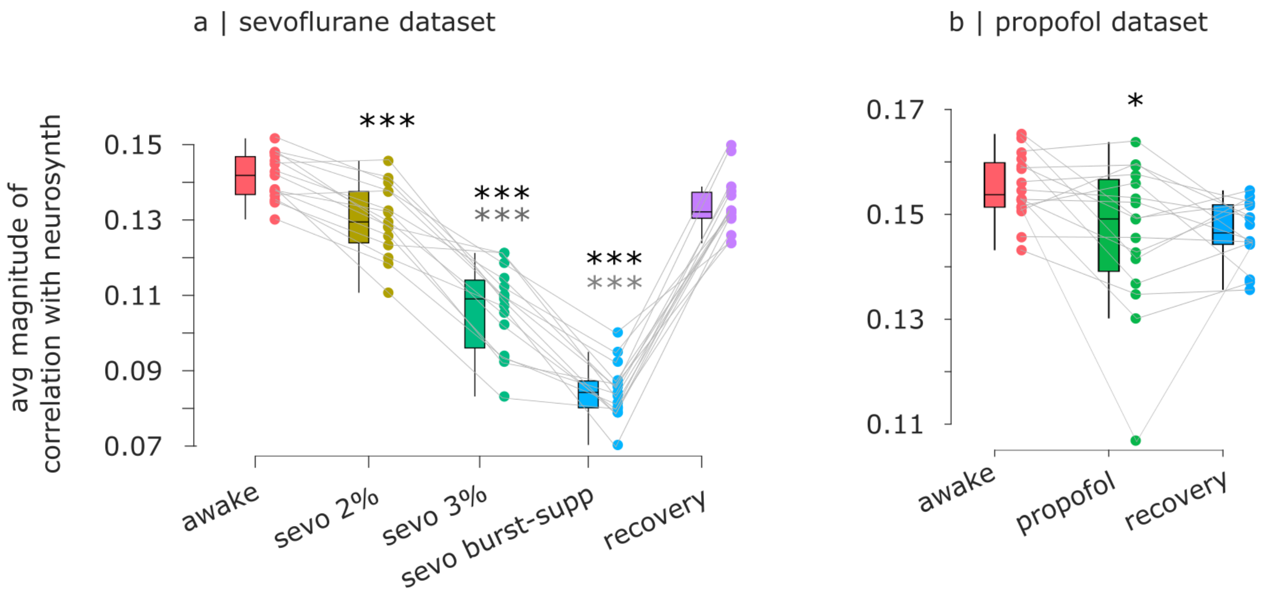


Figure S7. **Alternative quantification of cognitive matching from brain activity under wakefulness and anaesthesia |** (**a**) Sevoflurane dataset. (**b**) Propofol dataset. Ordinate: temporal average of the mean absolute value of spatial correlation between brain activity and 123 NeuroSynth meta-analytic maps. *** (black), $p < 0.001$ against wakefulness (FDR-corrected); ** (gray), $p < 0.001$ against recovery (FDR-corrected). Box-plot: center line, median; box limits, upper and lower quartiles; whiskers, $1.5\times$ interquartile range.

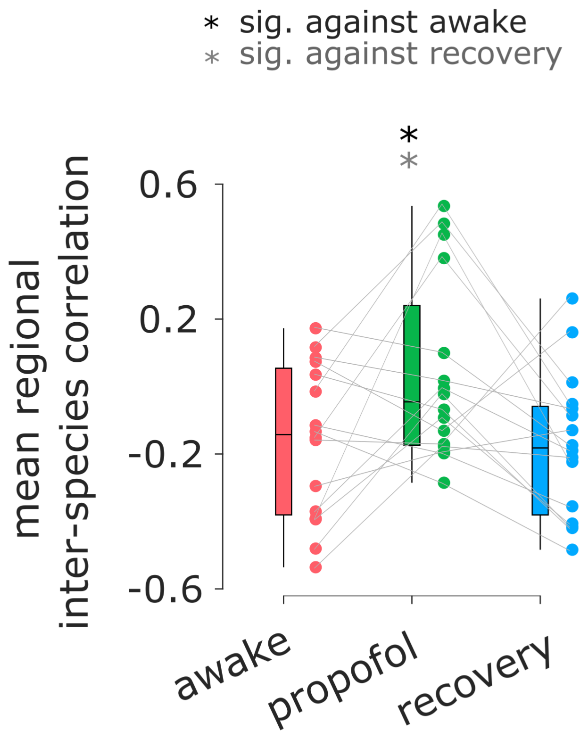


Figure S8. **Human cortical functional connectivity is more similar to macaque cortical connectivity under propofol anaesthesia than during wakefulness |** Ordinate: Regional correlation of mean FC between human and macaque across wakefulness, propofol anaesthesia, and recovery. * (black), $p < 0.05$ against wakefulness (FDR-corrected); * (gray), $p < 0.05$ against recovery (FDR-corrected). Box-plot: center line, median; box limits, upper and lower quartiles; whiskers, $1.5\times$ interquartile range.

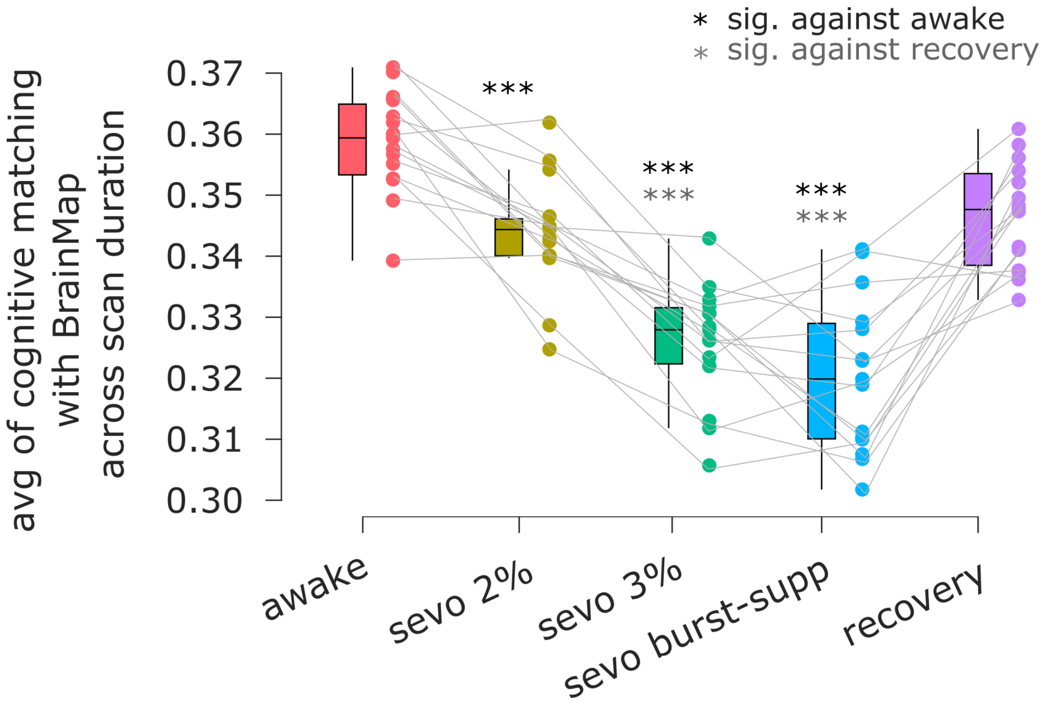


Figure S9. **Replication of decodability results using the BrainMap meta-analytic database |** Ordinate: mean across time of the best decoding score (maximum spatial correlation between brain activity and 66 meta-analytic maps from BrainMap). *** (black), $p < 0.001$ against wakefulness (FDR-corrected); *** (gray), $p < 0.001$ against recovery (FDR-corrected). Box-plot: center line, median; box limits, upper and lower quartiles; whiskers, $1.5\times$ interquartile range.

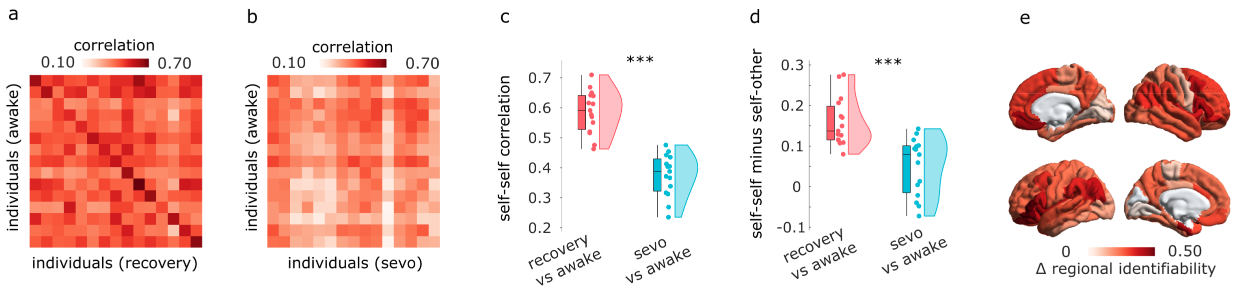


Figure S10. **Identifiability under anaesthesia is robust to parcellation choice |** (**a**) Identifiability matrix between wakefulness and post-anaesthetic recovery. (**b**) Identifiability matrix between wakefulness and sevoflurane anaesthesia (right). Entries along the diagonal, represent self-self similarity (correlation of FC patterns), whereas off-diagonal entries represent self-other similarity. (**c**) Self-self similarity is significantly higher between two conscious states, than between wakefulness and sevoflurane anaesthesia. (**d**) The difference between self-self correlation and mean self-other correlation (differential identifiability) is significantly higher between two conscious states, than between wakefulness and sevoflurane anaesthesia. (**e**) The regional distribution of contributions to identifiability (change in intra-class correlation coefficient) is plotted on the cortical surface for the 68 ROIs of the Desikan-Killiany atlas. Box-plot: center line, median; box limits, upper and lower quartiles; whiskers, $1.5\times$ interquartile range. ***, $p < 0.001$.

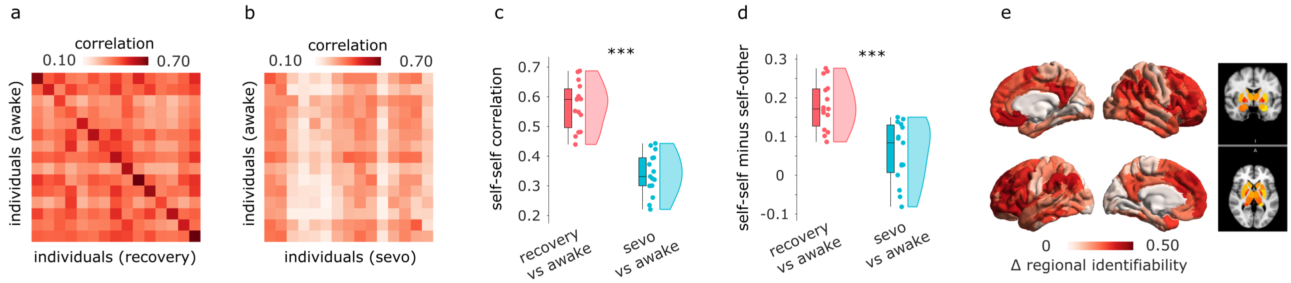


Figure S11. **Identifiability under anaesthesia is robust to inclusion of subcortex | (a)** Identifiability matrix between wakefulness and post-anaesthetic recovery. (**b**) Identifiability matrix between wakefulness and sevoflurane anaesthesia (right). Entries along the diagonal, represent self-self similarity (correlation of FC patterns), whereas off-diagonal entries represent self-other similarity. (**c**) Self-self similarity is significantly higher between two conscious states, than between wakefulness and sevoflurane anaesthesia. (**d**) The difference between self-self correlation and mean self-other correlation (differential identifiability) is significantly higher between two conscious states, than between wakefulness and sevoflurane anaesthesia. (**e**) The regional distribution of contributions to identifiability (change in intra-class correlation coefficient) is plotted on the cortical surface for the 200-ROIs of the Schaefer atlas, and plotted in volumetric space for the 32 ROIs of the Tian subcortical atlas. Box-plot: center line, median; box limits, upper and lower quartiles; whiskers, $1.5\times$ interquartile range. ***, $p < 0.001$.

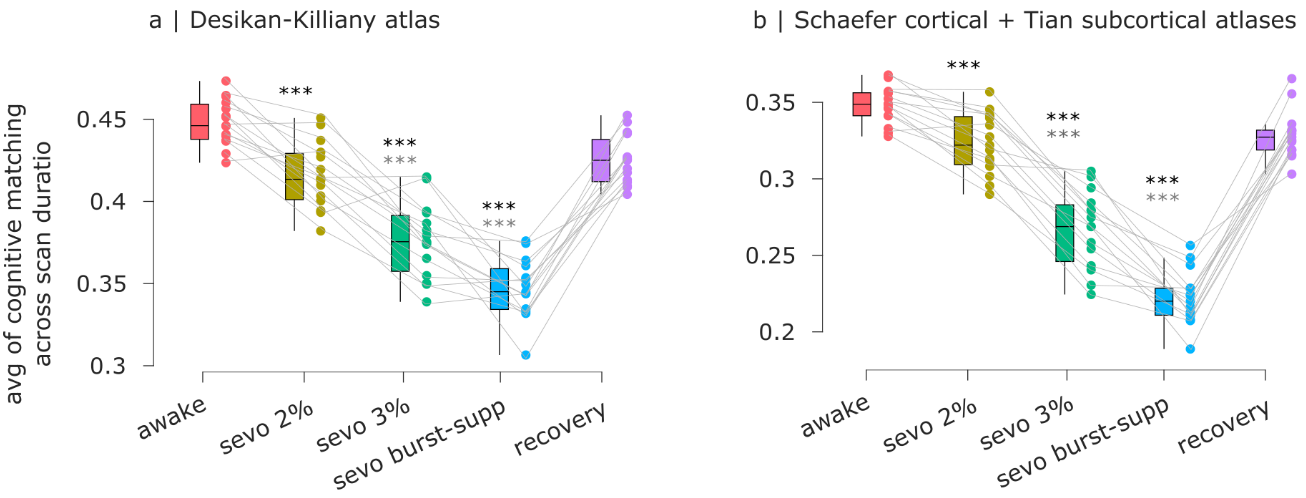


Figure S12. **Decodability of meta-analytic patterns is robust to parcellation choice and inclusion of subcortex | (a)** Results for Desikan-Killiany anatomical parcellation. (**b**) Results for the combined Schaefer-200 cortical atlas and Tian 32-ROI subcortical atlas. Ordinate: mean across time of the best decoding score (maximum spatial correlation between brain activity and 123 NeuroSynth meta-analytic maps). ***, $p < 0.001$ against wakefulness (FDR-corrected); *** (gray), $p < 0.001$ against recovery (FDR-corrected). Box-plot: center line, median; box limits, upper and lower quartiles; whiskers, $1.5\times$ interquartile range.

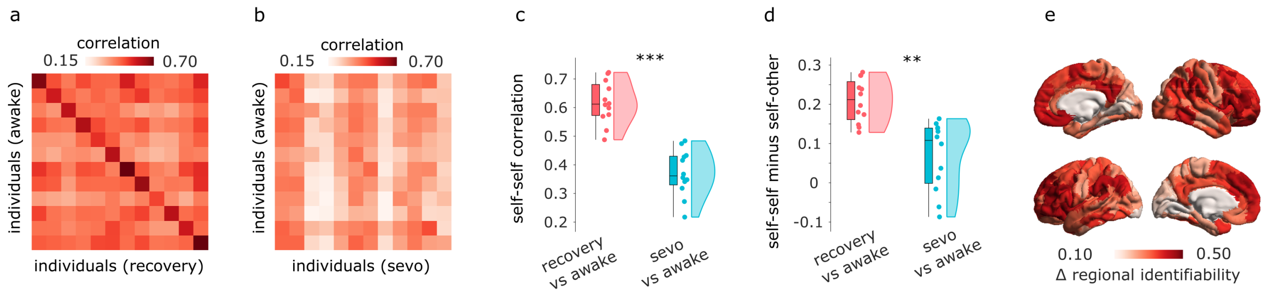


Figure S13. **Replication of identifiability results upon excluding high-motion individuals |** (**a**) Identifiability matrix between wakefulness and post-anaesthetic recoveryy. (**b**) Identifiability matrix between wakefulness and sevoflurane anaesthesia. Entries along the diagonal, represent self-self similarity (correlation of FC patterns), whereas off-diagonal entries represent self-other similarity. (**c**) Self-self similarity is significantly higher between two conscious states, than between wakefulness and sevoflurane. (**d**) The difference between self-self correlation and mean self-other correlation (differential identifiability) is significantly higher between two conscious states, than between wakefulness and sevoflurane. **, $p < 0.01$; ***, $p < 0.001$. (**e**) The regional distribution of contributions to identifiability (change in intra-class correlation coefficient) is plotted on the cortical surface. It is significantly spatially correlated with the corresponding map obtained when including all individuals: Spearman $\rho = 0.94$, $p_{spin} < 0.001$, $N = 200$ regions.

| Contrast | Baseline Mean | Baseline SD | Altered Mean | Null SD | tStat | df | pVal | Eff Size | 95% CI Lower | 95%CI Upper | Motion corr | Motion pVal |
|---|---|---|---|---|---|---|---|---|---|---|---|---|
| **Awake vs Vol 2%** | 3.81e-01 | 1.30e-02 | 3.54e-01 | 2.33e-02 | 5.60 | 14 | p <0.001 | 1.40 | 0.96 | 2.17 | -0.01 | 0.985 |
| **Awake vs Vol 3%** | 3.81e-01 | 1.30e-02 | 2.95e-01 | 2.90e-02 | 11.24 | 14 | p <0.001 | 3.68 | 2.91 | 5.38 | 0.23 | 0.404 |
| **Awake vs Burst-Supp** | 3.81e-01 | 1.30e-02 | 2.36e-01 | 1.68e-02 | 28.86 | 14 | p <0.001 | 9.28 | 7.35 | 13.93 | 0.27 | 0.327 |
| **Recovery vs Vol 2%** | 3.61e-01 | 1.81e-02 | 3.54e-01 | 2.33e-02 | 1.22 | 14 | 0.244 | 0.33 | -0.22 | 0.89 | -0.28 | 0.314 |
| **Recovery vs Vol 3%** | 3.61e-01 | 1.81e-02 | 2.95e-01 | 2.90e-02 | 8.38 | 14 | p <0.001 | 2.61 | 2.02 | 3.86 | 0.07 | 0.802 |
| **Recovery vs Burst-Supp** | 3.61e-01 | 1.81e-02 | 2.36e-01 | 1.68e-02 | 20.31 | 14 | p <0.001 | 6.85 | 5.44 | 10.56 | -0.11 | 0.695 |

TABLE S1. Statistical results for cognitive matching from NeuroSynth in the sevoflurane dataset.

| Contrast | Baseline Mean | Baseline SD | Altered Mean | Null SD | tStat | df | pVal | Eff Size | 95% CI Lower | 95%CI Upper | Motion corr | Motion pVal |
|---|---|---|---|---|---|---|---|---|---|---|---|---|
| **Awake vs Vol 2%** | -1.03e-01 | 2.24e-01 | 3.86e-03 | 2.64e-01 | -1.39 | 14 | 0.186 | -0.42 | -1.11 | 0.17 | 0.30 | 0.283 |
| **Awake vs Vol 3%** | -1.03e-01 | 2.24e-01 | 2.22e-01 | 2.42e-01 | -3.97 | 14 | 0.001 | -1.34 | -2.45 | -0.66 | 0.32 | 0.242 |
| **Awake vs Burst-Supp** | -1.03e-01 | 2.24e-01 | 4.86e-01 | 1.33e-01 | -11.16 | 14 | p <0.001 | -3.08 | -4.92 | -2.25 | -0.19 | 0.490 |
| **Recovery vs Vol 2%** | -1.31e-02 | 2.36e-01 | 3.86e-03 | 2.64e-01 | -0.19 | 14 | 0.850 | -0.07 | -0.76 | 0.63 | 0.64 | 0.012 |
| **Recovery vs Vol 3%** | -1.31e-02 | 2.36e-01 | 2.22e-01 | 2.42e-01 | -3.10 | 14 | 0.008 | -0.95 | -1.81 | -0.38 | 0.63 | 0.014 |
| **Recovery vs Burst-Supp** | -1.31e-02 | 2.36e-01 | 4.86e-01 | 1.33e-01 | -7.80 | 14 | p <0.001 | -2.51 | -4.10 | -1.70 | 0.60 | 0.020 |

TABLE S2. Statistical results for human-macaque functional similarity in the sevoflurane dataset.

| Contrast | Baseline Mean | Baseline SD | Altered Mean | Null SD | tStat | df | pVal | Eff Size | 95% CI Lower | 95%CI Upper | Motion corr | Motion pVal |
|---|---|---|---|---|---|---|---|---|---|---|---|---|
| **Awake vs Ppfl** | 4.13e-01 | 1.44e-02 | 3.96e-01 | 3.40e-02 | 2.03 | 15 | 0.061 | 0.63 | 0.10 | 1.17 | -0.61 | 0.014 |
| **Recovery vs Ppfl** | 3.93e-01 | 1.33e-02 | 3.96e-01 | 3.40e-02 | -0.38 | 15 | 0.710 | -0.11 | -0.96 | 0.41 | -0.72 | 0.002 |

TABLE S3. Statistical results for cognitive matching from NeuroSynth in the propofol dataset.

| Contrast | Baseline Mean | Baseline SD | Altered Mean | Null SD | tStat | df | pVal | Eff Size | 95% CI Lower | 95%CI Upper | Motion corr | Motion pVal |
|---|---|---|---|---|---|---|---|---|---|---|---|---|
| **Awake vs Ppfl** | -1.60e-01 | 2.28e-01 | 3.89e-02 | 2.71e-01 | -2.55 | 15 | 0.022 | -0.77 | -1.35 | -0.24 | -0.27 | 0.304 |
| **Recovery vs Ppfl** | -1.73e-01 | 2.13e-01 | 3.89e-02 | 2.71e-01 | -2.83 | 15 | 0.013 | -0.84 | -1.47 | -0.29 | -0.02 | 0.935 |

TABLE S4. Statistical results for human-macaque functional similarity in the propofol dataset

| Contrast | Baseline Mean | Baseline SD | Altered Mean | Null SD | tStat | df | pVal | Eff Size | 95% CI Lower | 95%CI Upper | Motion corr | Motion pVal |
|---|---|---|---|---|---|---|---|---|---|---|---|---|
| **Awake vs Vol 2%** | 4.48e-01 | 1.40e-02 | 4.16e-01 | 1.99e-02 | 5.97 | 14 | p <0.001 | 1.82 | 1.22 | 2.91 | 0.15 | 0.602 |
| **Awake vs Vol 3%** | 4.48e-01 | 1.40e-02 | 3.77e-01 | 2.24e-02 | 10.92 | 14 | p <0.001 | 3.69 | 2.85 | 5.60 | 0.32 | 0.248 |
| **Awake vs Burst-Supp** | 4.48e-01 | 1.40e-02 | 3.47e-01 | 1.81e-02 | 16.32 | 14 | p <0.001 | 6.03 | 4.81 | 9.13 | 0.26 | 0.340 |
| **Recovery vs Vol 2%** | 4.25e-01 | 1.51e-02 | 4.16e-01 | 1.99e-02 | 1.56 | 14 | 0.141 | 0.50 | -0.12 | 1.23 | -0.14 | 0.611 |
| **Recovery vs Vol 3%** | 4.25e-01 | 1.51e-02 | 3.77e-01 | 2.24e-02 | 7.71 | 14 | p <0.001 | 2.44 | 1.85 | 3.55 | 0.31 | 0.254 |
| **Recovery vs Burst-Supp** | 4.25e-01 | 1.51e-02 | 3.47e-01 | 1.81e-02 | 13.46 | 14 | p <0.001 | 4.51 | 3.58 | 6.66 | -0.02 | 0.954 |

TABLE S5. Statistical results for cognitive matching from NeuroSynth in the sevoflurane dataset, using the Desikan-Killiany anatomical atlas.

| Contrast | Baseline Mean | Baseline SD | Altered Mean | Null SD | tStat | df | pVal | Eff Size | 95% CI Lower | 95%CI Upper | Motion corr | Motion pVal |
|---|---|---|---|---|---|---|---|---|---|---|---|---|
| **Awake vs Vol 2%** | 3.59e-01 | 8.34e-03 | 3.44e-01 | 9.36e-03 | 5.28 | 14 | p <0.001 | 1.63 | 1.02 | 2.60 | 0.14 | 0.611 |
| **Awake vs Vol 3%** | 3.59e-01 | 8.34e-03 | 3.26e-01 | 9.65e-03 | 11.44 | 14 | p <0.001 | 3.50 | 2.70 | 5.16 | 0.41 | 0.130 |
| **Awake vs Burst-Supp** | 3.59e-01 | 8.34e-03 | 3.20e-01 | 1.26e-02 | 8.90 | 14 | p <0.001 | 3.44 | 2.51 | 5.47 | 0.17 | 0.549 |
| **Recovery vs Vol 2%** | 3.47e-01 | 8.65e-03 | 3.44e-01 | 9.36e-03 | 1.00 | 14 | 0.336 | 0.32 | -0.31 | 1.00 | -0.19 | 0.507 |
| **Recovery vs Vol 3%** | 3.47e-01 | 8.65e-03 | 3.26e-01 | 9.65e-03 | 7.06 | 14 | p <0.001 | 2.19 | 1.69 | 3.05 | 0.23 | 0.411 |
| **Recovery vs Burst-Supp** | 3.47e-01 | 8.65e-03 | 3.20e-01 | 1.26e-02 | 6.27 | 14 | p <0.001 | 2.34 | 1.61 | 3.75 | -0.09 | 0.763 |

TABLE S6. Statistical results for BrainMap decoding in the sevoflurane dataset

| Contrast | Baseline Mean | Baseline SD | Altered Mean | Null SD | tStat | df | pVal | Eff Size | 95% CI Lower | 95%CI Upper | Motion corr | Motion pVal |
|---|---|---|---|---|---|---|---|---|---|---|---|---|
| **Awake vs Vol 2%** | 3.48e-01 | 1.20e-02 | 3.23e-01 | 1.99e-02 | 5.73 | 14 | $p < 0.001$ | 1.45 | 0.99 | 2.24 | 0.26 | 0.340 |
| **Awake vs Vol 3%** | 3.48e-01 | 1.20e-02 | 2.67e-01 | 2.49e-02 | 12.93 | 14 | $p < 0.001$ | 3.98 | 3.22 | 5.69 | 0.25 | 0.375 |
| **Awake vs Burst-Supp** | 3.48e-01 | 1.20e-02 | 2.22e-01 | 1.76e-02 | 23.63 | 14 | $p < 0.001$ | 8.07 | 6.44 | 11.93 | 0.36 | 0.192 |
| **Recovery vs Vol 2%** | 3.29e-01 | 1.55e-02 | 3.23e-01 | 1.99e-02 | 1.03 | 14 | 0.318 | 0.29 | -0.29 | 0.85 | -0.11 | 0.705 |
| **Recovery vs Vol 3%** | 3.29e-01 | 1.55e-02 | 2.67e-01 | 2.49e-02 | 8.75 | 14 | $p < 0.001$ | 2.86 | 2.24 | 4.04 | 0.23 | 0.404 |
| **Recovery vs Burst-Supp** | 3.29e-01 | 1.55e-02 | 2.22e-01 | 1.76e-02 | 18.81 | 14 | $p < 0.001$ | 6.20 | 4.69 | 10.51 | 0.10 | 0.714 |

TABLE S7. Statistical results for cognitive matching from NeuroSynth in the sevoflurane dataset, using 200 cortical regions from the Schaefer atlas, and 32 subcortical regions from the Tian atlas.
